# Sexual signals persist over deep time: ancient co-option of bioluminescence for courtship displays in cypridinid ostracods

**DOI:** 10.1101/2021.09.03.458903

**Authors:** Emily A. Ellis, Jessica A. Goodheart, Nicholai M. Hensley, Vanessa L. González, Nicholas J. Reda, Trevor J. Rivers, James G. Morin, Elizabeth Torres, Gretchen A. Gerrish, Todd H. Oakley

**Author notes:** Equal authorship (Emily A. Ellis, Jessica A. Goodheart).

## Abstract

Although the the diversity, beauty, and intricacy of sexually selected courtship displays command the attention of evolutionists, the longevity of these traits in deep time is poorly understood. Population-based theory suggests sexual selection could either lower or raise extinction risk, resulting in high or low persistence of lineages with sexually selected traits. Furthermore, empirical studies that directly estimate longevity of sexually selected traits are uncommon. Sexually selected signals - including bioluminescent courtship - originated multiple times during evolution, allowing empirical study of their longevity after careful phylogenetic and divergence time analyses. Here, we estimate the first transcriptome-based molecular phylogeny and divergence times of Cypridinidae. We report extreme longevity of bioluminescent courtship, a trait important in mate choice and probably under sexual selection. Our relaxed-clock estimates of divergence times coupled with stochastic character mapping show luminous courtship evolved only once in Cypridinidae in a Sub-Tribe we name Luxorina at least 151 Million Years Ago (Ma) from cypridinid ancestors that used bioluminescence only in anti-predator displays, defining a Tribe we name Luminini. This time-calibrated molecular phylogeny of cypridinids will serve as a foundation for integrative and comparative studies on the biochemistry, molecular evolution, courtship, diversification, and ecology of cypridinid bioluminescence. The persistence of luminous courtship for hundreds of millions of years indicates that rates of speciation within the group exceeded extinction risk, allowing for the persistence of a diverse clade of signalling species and that sexual selection did not cause rapid loss of associated traits.

## Introduction

By producing some of the most diverse, beautiful, and elaborate biological displays, the evolution of sexually selected traits is a topic of fundamental importance. While sexually selected traits in a handful of model clades (Prum 2017; Ryan et al. 2019) are the focus of much study, open questions remain, such as the longevity of such traits (Wiens and Tuschhoff 2020). Predicting longevity during macroevolution of sexually selected traits is difficult because population-based inferences that relate sexual selection to extinction are mixed (Kokko and Brooks 2003; Martínez-Ruiz and Knell 2017). On one hand, sexual selection could reduce mutation load, increase mean fitness, and decrease extinction (Tomkins et al. 2004; Long et al. 2012; Plesnar-Bielak et al. 2012; Lumley et al. 2015). On the other hand, costs imposed by sexual selection - through maladaptation of exaggerated traits and/or reduced effective population - could decrease population fitness and increase extinction (Promislow 1992; Tanaka 1996; Sorci et al. 1998; Doherty et al. 2003; Morrow and Fricke 2004). Together, these studies show many factors - especially population size, environmental variability, honesty of signal, and eco-evolutionary feedbacks - dictate relationships between sexual selection and extinction risk at population scales. If these factors vary across species during macroevolutionary radiations, the longevity of a sexually selected trait will not be predicatable for individual populations or species. In addition, few empirical studies of trait duration are available, although examples of ancient signals, including those with modern sexual functions, date to at least 200 Ma. Examples include acoustic signals in tetrapods (Chen and Wiens 2020) and insects (Gu et al. 2012; Song et al. 2020), luminous courtship in terrestrial fireflies (Powell et al.; Höhna et al. 2021), and the role of flowers in plant-pollinator interactions (Bao et al. 2019; van der Kooi and Ollerton 2020). Conversely, extinction rates increased in fossil ostracods that are sexually dimorphic in body size, where the dimorphism may be a strong proxy for post-copulatory sexual selection (Martins et al. 2018). Therefore, the longevity of sexually selected traits remains largely untested, despite implications for understanding the power of sexual selection to influence biodiversity on macroevolutionary timescales.

To study the origin and longevity of sexually selected traits in macroevolution, a valuable trait is bioluminescence, which evolved separately at least 94 times (Lau and Oakley 2020), providing replicative power for comparative analyses (Ellis and Oakley 2016). Bioluminescence has multiple organismal functions including defense and reproduction, which affect ecosystems (Morin 1983; Wilson and Hastings 1998; Haddock et al. 2010; Widder 2010). When used for courtship, bioluminescence is associated with more species compared to sister clades that lack luminous courtship, providing an example of a sexually selected trait that may increase rates of speciation (Ellis and Oakley 2016). Unfortunately, the specific number and timing of origins of bioluminescence remain uncertain for luminous groups and their relatives, despite some recent attention (see Ellis and Oakley 2016 for summary of ages). Here, we report new phylotranscriptomic analyses and divergence time estimates to understand the origins and longevity of bioluminescence and luminous courtship in cypridinid ostracods, a family of crustaceans that contains at least 250 species across 30 genera (Morin 2019), and which evolved bioluminescence separately from other animals.

Bioluminescent courtship in cypridinids is almost certainly influenced by sexual selection, as information rich, sexually dimorphic traits that allow for mate choice (Andersson 1994; Rivers and Morin 2013). While all known bioluminescent cypridinids can produce anti-predatory displays by secreting defensive clouds of light in response to predation attempts (Morin and Cohen 2010; Rivers and Morin 2012; Morin 2019), only about 75 described and nominal species, restricted to the Caribbean, also produce bioluminescent courtship signals (Morin 1986, 2019; Morin and Cohen 2010; Reda et al. 2019). Beginning near twilight’s end (Gerrish et al. 2009), often in multi-species assemblages of 6-8 species that overlap in time and space (Gerrish and Morin 2016), male cypridinids produce patterns of ephemeral pulses of light, secreted while swimming in tight spirals (Rivers and Morin 2008). Like other radiations of sexually selected signals (Prum 2017), species-specific displays in signaling cypridinids vary considerably in many parameters (Morin 1986, 2019; Gerrish et al. 2009; Gerrish and Morin 2016; Hensley et al. 2019, 2021).

Previous phylogenetic analyses of morphology (Cohen and Morin 2003) and rDNA (Torres and Gonzalez 2007) both indicated a single origin of bioluminescence and suggested a single origin of luminous courtship in Cypridinidae (Morin 2019). However, beyond the topology itself, we have very little information about divergence times for these evolutionary transitions. One study of divergence times of Pancrustacea did include two luminous ostracod species, but resulted in very large confidence intervals on the time of origin of their common ancestor, ranging from about 50-250 Ma (Oakley et al. 2013). Here, we present the first transcriptomic phylogenies and divergence time estimates for Cypridinidae, which indicate an ancient origin of bioluminescence and a long persistence of luminous courtship, a trait almost certainly under sexual selection (Morin and Cohen 2010). We conservatively estimate the origin of bioluminescent courtship in cypridinids to be at least 151 Ma. If researchers continue to identify long persistence of traits under sexual selection, it would support these traits as often associated with increases in the rate of speciation relative to their effect on extinction.

## Materials and Methods

### Sampling Strategy and Collection

Based on previous morphological phylogenies and known distributions of species, we aimed to balance cost-efficiency and diversity of sampling by choosing eight primary sampling locations. We targeted species with luminous courtship mainly from five localities: Jamaica, Honduras (Roatan), Belize, Panama, and Puerto Rico. We targeted species outside the courtship signaling clade from three localities: Australia, Japan, and the United States. We collected ostracods via net sampling during courtship displays, sediment sampling, or carrion traps (see Cohen and Oakley (2017) for sampling methods and Table S1 for taxa sampled). We preserved samples in 95% ethanol for vouchers and RNAlater for sequencing. To collect taxa with luminous courtship (where many species are undescribed), we targeted unique courtship displays to isolate individual species because ecologically overlapping courtship signals are distinct enough to distinguish species (Gerrish and Morin 2016). We also measured length and height of carapaces to assign species to genera in the field, prior to molecular analysis, and to affirm single-species identity of pooled individuals (Table S2). The ratio of carapace length to height is usually diagnostic for genera of signaling cypridinids and distinguishes sympatric species in combination with signals (Gerrish and Morin 2016; Hensley et al. 2019; Reda et al. 2019).

### Taxon Sampling and Sequencing

We generated new RNA-Seq data from 45 species of cypridinid ostracods and four species of non-cypridinid ostracods. We included outgroups from all four non-cypridinid families of Myodocopida, sequencing one species each of Rutidermatidae and Sarsiellidae, and two of Philomedidae. We also used published sequence data from a cylindroleberid (SRX2085850) (Schwentner et al. 2018), and predicted proteins from the genome of *Darwinula stevensoni* (ENA PRJEB38362) (Schön et al. 2021). For transcriptome sequencing, we combined RNA from whole bodies of up to 10 pooled individuals (collected by signal but with identity corroborated by morphological features). Pooling is necessary due to the small size of the animals (≤ 2 mm) and low RNA yield. For some samples we extracted RNA using Trizol, and for the rest we used Qiagen RNEasy Kits. We either used the Illumina or NEBNext Ultra RNA Library Prep Kits to prepare transcriptome libraries (Table S1).

### Quality Control and Dataset Assembly

We trimmed RNA-seq adapters and low-quality bases (<20) using TrimGalore v0.4.1 (Krueger 2012), then assembled trimmed reads using Trinity v2.2.0 (Grabherr et al. 2011). In cases of multiple transcriptomes from the same species, we combined raw, trimmed reads to create single assemblies. We used CroCo v0.1 (Simion et al. 2018) to remove contigs in a given assembly both found and expressed at a higher level in a transcriptome from another species (that was sequenced on the same lane), which implies cross-contamination. We assessed completeness using BUSCO v3.0.1 (Simão et al. 2015) using the arthropod_odb9 single-copy ortholog database. We generally required transcriptomes to have at least 100 complete BUSCO genes for further analyses. However, we made exceptions for three species, (*Photeros jamescasei, Jimmorinia gunnari*, and *Melavargula japonica*), to maintain these described and taxonomically important species within the analyses. We produced protein predictions using TransDecoder v3.0.1 (Haas et al. 2013), with default parameters.

### Orthology Determination

For analyses that relied on comparing orthologs (excluding paralogs), we used Orthofinder (Emms and Kelly 2019) plus phylopypruner. We grouped genes with Orthofinder 2.5.2 (Emms and Kelly 2019), using an MCL inflation parameter set to 2.1 and used Diamond as the similarity search program. This allowed us to cluster sequences based on similarity into “orthogroups” that contain a mix of orthologs and paralogs. Next, we created gene trees for the separate orthogroups by aligning each using the ‘auto’ method of MAFFT v7.305 (Katoh and Standley 2013) and using maximum likelihood implemented in FastTree v2.1.9 (Price et al. 2010), chosen for its speed and displayed in final results because are congruent with concatenation methods using IQ-TREE2 (Minh et al. 2020), and assuming a JTT+CAT. We used phylopypruner 1.2.3 (Thalén 2018) to remove paralogs, requiring a minimum of 20 species for each gene family, minimum support values of 0.7, and used the LS pruning method. We refer to this resulting set of orthologs as the “ppp orthologs”.

### Previously Published Datasets

We also added single-gene mitochondrial data (12S, 16S, CO1), summarized in Table S5 reported previously in conference proceedings (Torres and Gonzalez 2007) or publications (Ogoh and Ohmiya 2004; Wakayama and Abe 2006; Wang et al. 2019; Goodheart et al. 2020; Pham et al. 2020). We then identified these three mitochondrial genes from transcriptomes by finding the longest contigs with high similarity to published complete mitochondrial genomes of *V. hilgendorfii* and *V. tsujii* (Ogoh and Ohmiya 2004; Goodheart et al. 2020). We annotated mitochondrial data from transcriptomes using MITOS (Bernt et al. 2013), then aligned each mitochondrial gene using ‘auto’ in MAFFT v7.305 (Katoh and Standley 2013).

### Phylogenetic Reconstruction

We compared coalescence and concatenation-based analyses using Maximum Likelihood and Bayesian approaches. For coalescence-based approaches, we used ASTRAL-Pro (Zhang et al. 2020), which assumes the multi-species coalescent to estimate a species tree from multiple gene trees, including paralogs. We used the same gene trees as input for ASTRAL-Pro as phylopypruner (described above). Next, we determined the best-fit partitioning scheme for each concatenated dataset using PartitionFinder v.2.1.1 (Lanfear et al. 2016) with RAxML (Stamatakis 2014) and rcluster (Lanfear et al. 2014). Using optimal partitioning schemes and models, we produced species trees in IQ-TREE v2.0.3, with additional options for -bnni and -alrt (set to 1000). We calculated gene Concordance Factors (gCF) using IQ-TREE (Minh et al. 2020).

### Divergence time estimation

We estimated divergence times of nodes in the cypridinid phylogeny using a Bayesian relaxed molecular clock approach in MrBayes (Ronquist and Huelsenbeck 2003), which offers a diversity of models, estimates tree topology and branch lengths in a Bayesian framework, and has a scriptable command line interface, leading to more repeatable science. Because Bayesian methods are computationally demanding (Smith et al. 2018), we chose 15 orthologs to use for divergence time estimation, as follows. We used sortadate (Smith et al. 2018) to rank all gene trees based on their consistency with the species tree inferred from all genes in ASTRAL-Pro and based on the most consistent root-to-tip branch lengths. To test the sensitivity of divergence time estimates to the number of genes selected, we compared results from 15 gene trees to age estimates from 20, 25, and 30 gene trees (Fig. S10). Although ostracods have a dense fossil record, most fossil ostracods are podocopids and assigning fossils confidently to taxonomic groups is often challenging (Tinn and Oakley 2008), restricting us to two fossil constraints. The first was Myodocopida, for which we used an offset exponential prior with a minimum age of 448.8 Ma and maximum of 509 Ma (Oakley et al. 2013; Wolfe et al. 2016). We also used a uniform prior between 443.8 and 509 Ma on the root of the tree, which is the common ancestor of Ostracoda. Along with node constraints, we used the Independent Gamma Rate (IGR) model (Lepage et al. 2007; Zhang 2016). We used a prior of exp(10) for variation in the IGR model. For the prior on the overall rate of the molecular clock (substitution rate/site/Myr), we assumed a lognormal distribution with a mean of 0.001 and standard deviation of 0.0007. For the model of amino acid substitution, we assumed fixed-rate models and used the command aamodelpr=mixed, which explores multiple fixed-rate models in proportion to their posterior probabilities. With these models and priors, we ran 500K steps of Markov Chain Monte Carlo (MCMC) in two chains, using a starting phylogeny based on parsimony.

### Ancestral state reconstructions

We reconstructed ancestral states for bioluminescence (presence/absence) and bioluminescent courtship (presence/absence) using our tree generated with ASTRAL-Pro. In order to create an ultrametric tree for ancestral state reconstruction analysis, we used mean ages for nodes matched to those from our Bayesian relaxed clock analysis. We set matching branch lengths using the ph_bladj function in the R package phylocomr (Webb et al. 2008; Ooms and Chamberlain 2019). We assessed fit for two models (for each character) using corrected Akaike Information Criterion (AICc), where: (i) transition rates were equal (ER), and (ii) forward and reverse transitions were different between states (all rates different, ARD). The ER model (AICc = 24.02231) was slightly better than the ARD model (AICc = 25.72293) for bioluminescence and for bioluminescent courtship (ER AICc = 23.799; ARD AICc = 24.70163), so we used ER for final ancestral state reconstruction analyses in R using the rayDISC function in corHMM (Beaulieu et al. 2013). To test for significance of the reconstruction at each node, we used proportional likelihood significance tests assuming a log likelihood difference of 2 or greater represents a significant difference (Pagel 1999).

To estimate timing of cypridinid bioluminescence and luminous courtship, we conducted time-tree stochastic character mapping (ttscm) (Alexandrou et al. 2013), which simulates character evolution with stochastic character mapping (Huelsenbeck et al. 2003) along time-calibrated phylogenies from the MCMC search to represent the posterior distribution of trees. While ancestral state reconstructions depict character state changes only at nodes, stochastic character maps allow state-changes along branches. When branches are time-calibrated, this approach provides absolute estimates of the timing of character state changes. We used stochastic character mapping in phytools (Revell 2012) using the same character states and models as described for ancestral state reconstruction. We used 100 time-calibrated trees from the MCMC search to represent the posterior probability distribution of tree topologies and branch lengths, and summarized the timing character state changes in histograms.

## Results

### Assembly and orthology statistics

Our final matrices of transcriptomic data (Table S3) include 45 cypridinids from 15 of 25 genera, plus six outgroup taxa representing all four other myodocopid families and the previously published genome sequence of one podocopid (Schön et al. 2021). Some of these transcriptomes were used previously to identify luciferase genes (Hensley et al. 2021).

### Evolution of luminous courtship and cypridinid bioluminescence

Our phylogenetic results support a single clade containing all Cypridinidae that use bioluminescence for courtship signaling (Figs 1, 3; Table 1). For this clade, we propose the Sub-Tribe name Luxorina (lux= light, uxoriae ~= courtship). Although an important consideration for defining Luxorina, our molecular data unfortunately does not contain the monotypic genus *Enewton* (Cohen and Morin 2010). Despite returning to the same collection site in Jamaica as Cohen and Morin (2010), we could not find *E. harveyi* (due to inclement weather). In previous parsimony analyses of morphological traits (Cohen and Morin 2003, 2010), *E. harveyi* is the sister of the rest of Luxorina. In the absence of molecular data, *Enewton* as sister to the rest of Luxorina remains the best hypothesis and we propose to include *Enewton* taxonomically within Luxorina.

**Figure 1.**
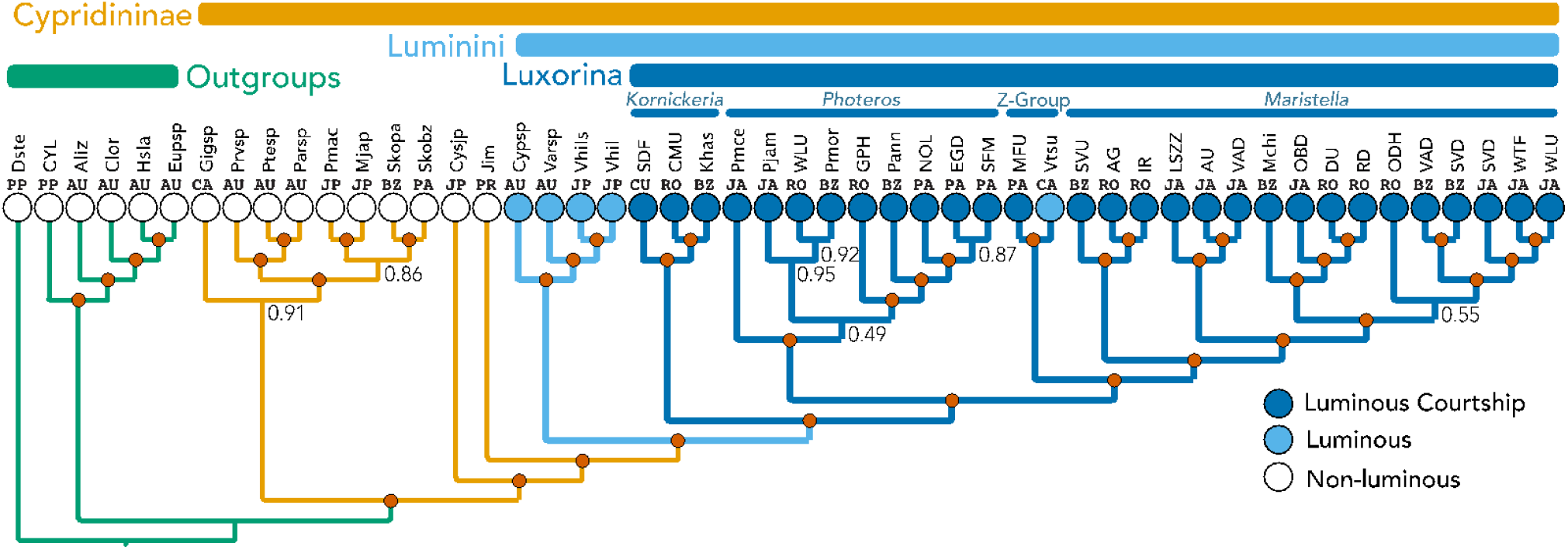
Species phylogeny inferred using multi-species-coalescent and transcriptome data. Red circles at nodes indicate 1.0 Posterior Probability, and nodes with values less than 1.0 are indicated numerically. Circles at tips represent non-luminous (white), luminous but non-courtship signaling (light blue), and species with luminous courtship (dark blue). The three described genera of signaling species are labeled at the top, along with “Z-group”, which is a clade of signaling species plus *Vargula tsujii. Vargula* is polytypic (see also (Cohen and Morin 2003; Morin 2019)) and the type species is outside this group, so formally, these should not be *Vargula*. Above the tree, we also labeled non-cypridinid Outgroups (green), and Cypridinidinae (Orange). We herein suggest a name for the bioluminescent Cypridinidae to be Tribe Luminini (for lumen = light and the typical Tribe suffix for zoology) and a name for the clade containing courtship-signaling cypridinids to be Sub-Tribe Luxorina (lux = light + uxoriae ~= courtship + -ini = typical zoological subtribe suffix). Locality abbreviations for collecting sites of species are above each tip as follows: AU = Australia, BZ = Belize, CA = California, USA, CU = Curacao, JA = Jamaica, JP = Japan, PA = Panama, PP = Previously Published, PR = Puerto Rico, and RO = Roatan, Honduras (see Table S1 for specific localities). Above locality names are abbreviations for species. Described species are abbreviated as the first letter of genus and first three letters of species epithet; undescribed species outside of Luxorina are labeled as first three letters of genus followed by sp or two locality or collector letters when there is more than one undescribed species from the genus; within Luxorina, we used 2-4 letter field codes for undescribed species (see supplement for tree with full species names).

**Table 1.**
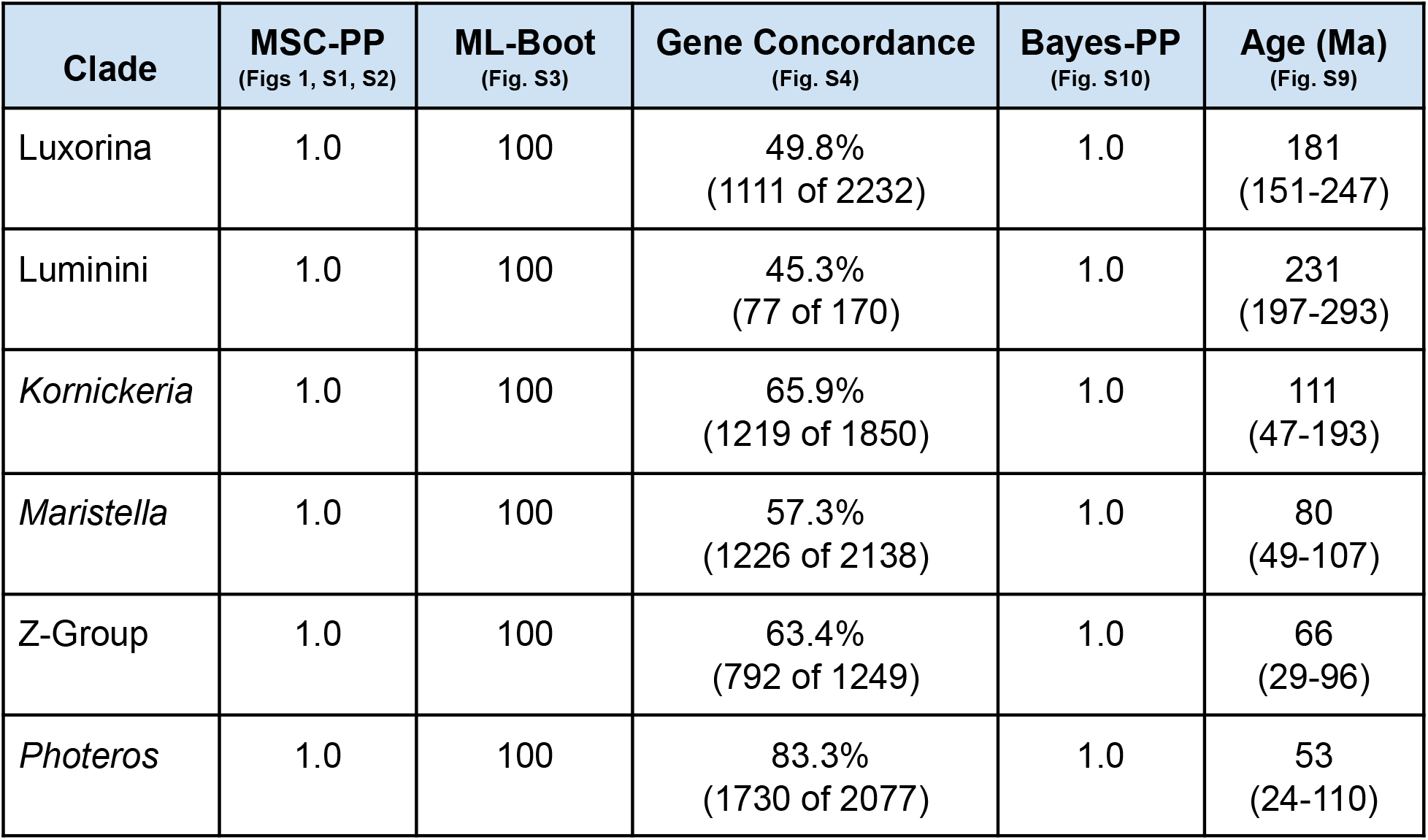
Summary of support for clades of interest from transcriptome data.

| **Clade** | **MSC-PP**<br>(Figs 1, S1, S2) | **ML-Boot**<br>(Fig. S3) | **Gene Concordance**<br>(Fig. S4) | **Bayes-PP**<br>(Fig. S10) | **Age (Ma)**<br>(Fig. S9) |
| --- | --- | --- | --- | --- | --- |
| Luxorina | 1.0 | 100 | 49.8%<br>(1111 of 2232) | 1.0 | 181<br>(151-247) |
| Luminini | 1.0 | 100 | 45.3%<br>(77 of 170) | 1.0 | 231<br>(197-293) |
| *Kornickeria* | 1.0 | 100 | 65.9%<br>(1219 of 1850) | 1.0 | 111<br>(47-193) |
| *Maristella* | 1.0 | 100 | 57.3%<br>(1226 of 2138) | 1.0 | 80<br>(49-107) |
| Z-Group | 1.0 | 100 | 63.4%<br>(792 of 1249) | 1.0 | 66<br>(29-96) |
| *Photeros* | 1.0 | 100 | 83.3%<br>(1730 of 2077) | 1.0 | 53<br>(24-110) |

Our results also support a single clade containing all bioluminescent Cypridinidae (Table 1). For this clade of bioluminescent cypridindids, and noting this is a separate origin of bioluminescence from halocyprid ostracods (Oakley 2005) that also involves a different chemistry (Campbell and Herring 1990), we propose the name Luminini (lumen = light). While supported in each of our analyses, the taxa available vary across data sets due to challenges obtaining transcriptomic data from some species. In the transcriptome-based analyses, bioluminescent cypridinids share a common ancestor in this clade with only luminous species. The dataset that combines mitochondrial and transcriptome data also supports Luminini (Fig. 3). However, the difference in taxon sampling highlights an important result. Namely, four species (three luminous and one unknown) with only mitochondrial data form a rather weakly supported clade (0.6 PP) with the non-luminous *Jimmorinia*, which is the sister group of luminous taxa in the transcriptome analyses. The three luminous species in this clade include *Vargula norvegica* from the deep Atlantic Ocean, and *Sheina orri* and *Vargula karamu* from Australia. *Vargula tubulata* is also in this clade, but the status of bioluminescence in the species is unknown. These species may form a clade within Luminini that also includes non-luminous *Jimmorinia*, leading to an inference of loss of bioluminescence in that genus (Figs 3, S12).

### Genus-level relationships within Cypridinidae

Our analysis included a number of undescribed species that we first assigned to genera using length to height ratio of the carapaces in the field (Table S2). Our phylogenetic reconstructions (Fig. 1) based on genetic data provide good support for these initial generic designations. Using these designations, both concatenated (Fig. S3) and coalescent approaches (Fig. 1) recovered four monophyletic genera (*Kornickeria, Photeros*, and *Maristella* and the not yet described “Z-Group”), all with strong support (Table 1). Additionally, *Maristella* (Reda et al. 2019), includes two distinct monophyletic clades, the small-bodied H-Group, which would remain in *Maristella*, and the larger-bodied U-Group (BZ-SVU, RO-AG, and RO-IR), which could be a new genus (Cohen and Morin 1989) (Fig. 1).

Furthermore, while strongly supported by transcriptome data, results of *Maristella* deserve more discussion because two other species (*Vargula contragula* and an undescribed species from Florida with only mitochondrial data) are closely related to *Maristella*, and may belong to yet another (undescribed) genus, provisionally called the C-group (Cohen and Morin 1986; Morin and Cohen 2017). Our results support the *Maristella* clade with high probability (1.0) to include both C-group and *Maristella*, but the position of the two C-group species compared to *Maristella* is unresolved (Fig. 3, S10). Confidently resolving relationships of groups similar, but perhaps sister to *Maristella* will require more data.

Another group within Luxorina includes representatives of what is provisionally called “Z-group” (Cohen and Morin 1986; Morin and Cohen 1988, 2017), which will need formal description as a new genus. Our analyses that add taxa with only mitochondrial data include three Panamanian species in a clade with *V. tsujii*: undescribed MFU, *“Vargula” kuna*, and *“Vargula” mizonoma*. The latter two were previously called Z-group (Morin and Cohen 1988; Cohen and Morin 2003). Therefore, we report strong support for a clade that includes the Z-group and *Vargula tsujii*. Even though *V. tsujii* is a described species, the genus should not be named *Vargula* (Cohen and Morin 2003) because the type species is the distantly related *V. norvegica* (Fig. 3). These results reinforce the highly polyphyletic nature of *Vargula* (Cohen and Morin 2003) and uncover the closest relatives of the only Luxorina species known from the Pacific, *V. tsujii*.

We also find strong support (1.0) for the relationships among courtship signaling genera, with *Kornickeria* sister to a clade containing *Photeros, Maristella*, Z-Group, and C-group (Fig. 3). Next, *Photeros* is the sister clade (0.76) to *Maristella*, Z-Group, and C-group, with Z-group as the sister of *Maristella* plus C-group (1.0). The relationships among these genera in Luxorina varied among previous studies.

### Divergence Time Estimates

Relaxed molecular clock analyses place the origin of the clade of species with bioluminescent courtship between (=Luxorina) 151-248 Ma (Bayesian Highest Posterior Density) with a median value of 197 Ma (Fig. 2A). Using time tree stochastic character mapping (ttscm) (Alexandrou et al. 2013), we estimate the character state transition to luminous courtship occurred 213 Ma (Fig. 2C). The estimates for the origin of cypridinid bioluminescence (=Luminini) are less clear because of issues with taxon sampling and the low quality of the transcriptome of *Jimmorinia*. Namely, the data set used for divergence times does not contain any species of the clade containing *Jimmorinia, Vargula norvegica*, or the other luminous Australian species; these taxa form the sister group to the rest of the luminous cypridinids (Fig. 1). Nevertheless, we can estimate the age of the common ancestor of all luminous species with transcriptome data as a minimum estimate for the origin of cypridinid bioluminescence to be 197-293 Ma with a median value of 237 Ma. Similarly, we can estimate the age of the common ancestor of luminous cypridinids and their closest non-luminous relative as a maximum age to be 233-361 Ma with a median value of 300 Ma (Fig. 2A). Using time tree stochastic character mapping (ttscm) (Alexandrou et al. 2013), we estimate the origin of bioluminescence occurred 267 Ma (Fig. 2B). We also note our estimates of even the most closely related species are always more than 8 Ma (Fig. S9), even for two populations of *V. hilgendorfii*. Although populations of *V. hilgendorfii* are quite diverged (Ogoh and Ohmiya 2005) our estimates seem very old at the tips of the tree, and we speculate this could be an artifact of extrapolating young nodes from the deepest nodes in the absence of other suitable fossils.

**Figure 2.**
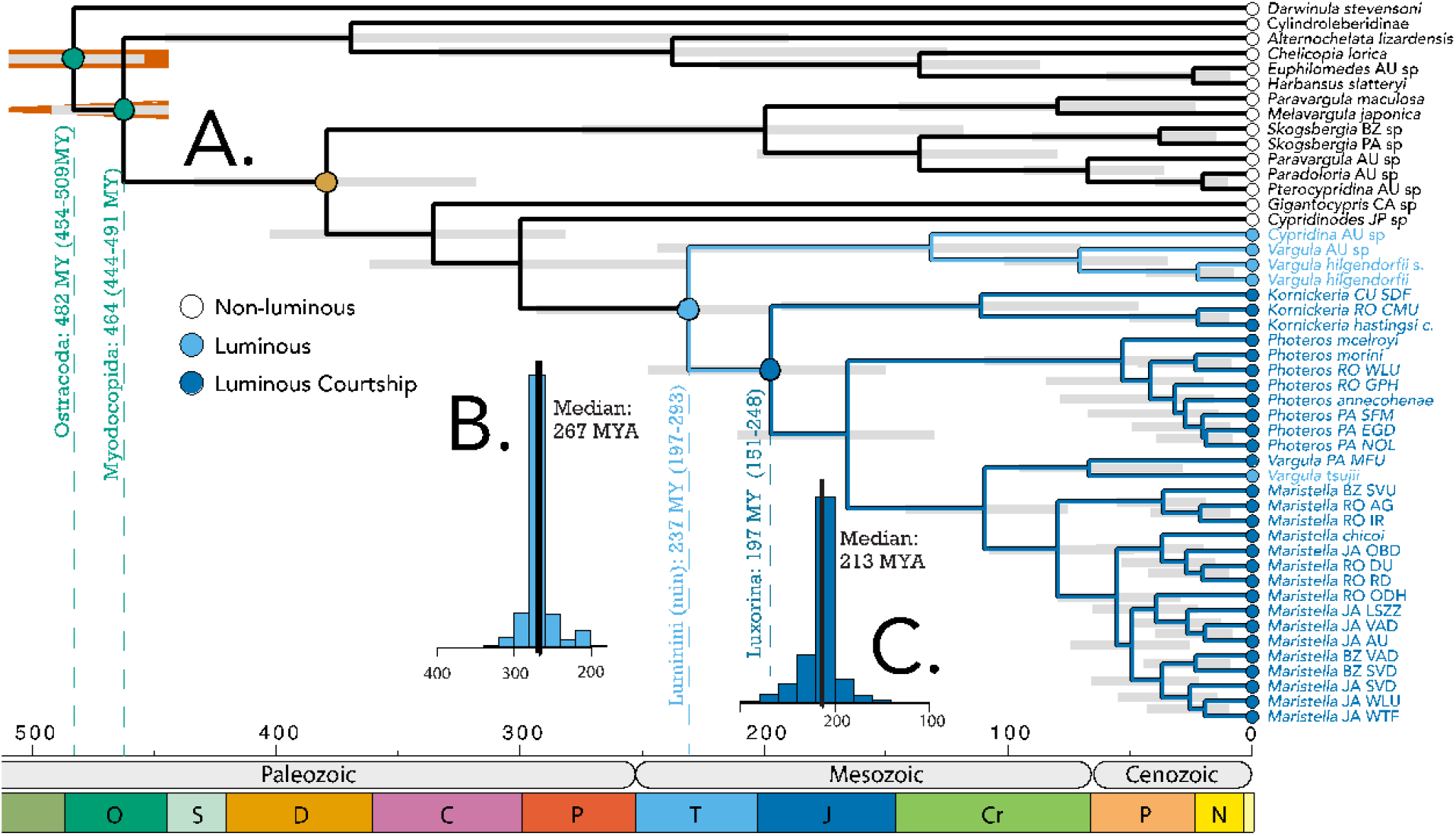
Time of taxon origins based on relaxed molecular clock analysis. **A.** We used two fossil constraints (green circles): Myodocopida, with offset exponential prior with a minimum of 448.8 Ma and maximum of 509 Ma and Ostracoda with a uniform prior between 443.8 and 509 Ma (Oakley et al. 2013; Wolfe et al. 2016) to estimate divergence times across our phylogeny; **B.** Results of time-tree stochastic character mapping (ttscm) estimates origin of bioluminescence to be 267 Ma because that is the median age of character state transitions to bioluminescence; **C**. Similarly, ttscm estimates the origin of luminous courtship to be 213 Ma.

**Figure 3.**
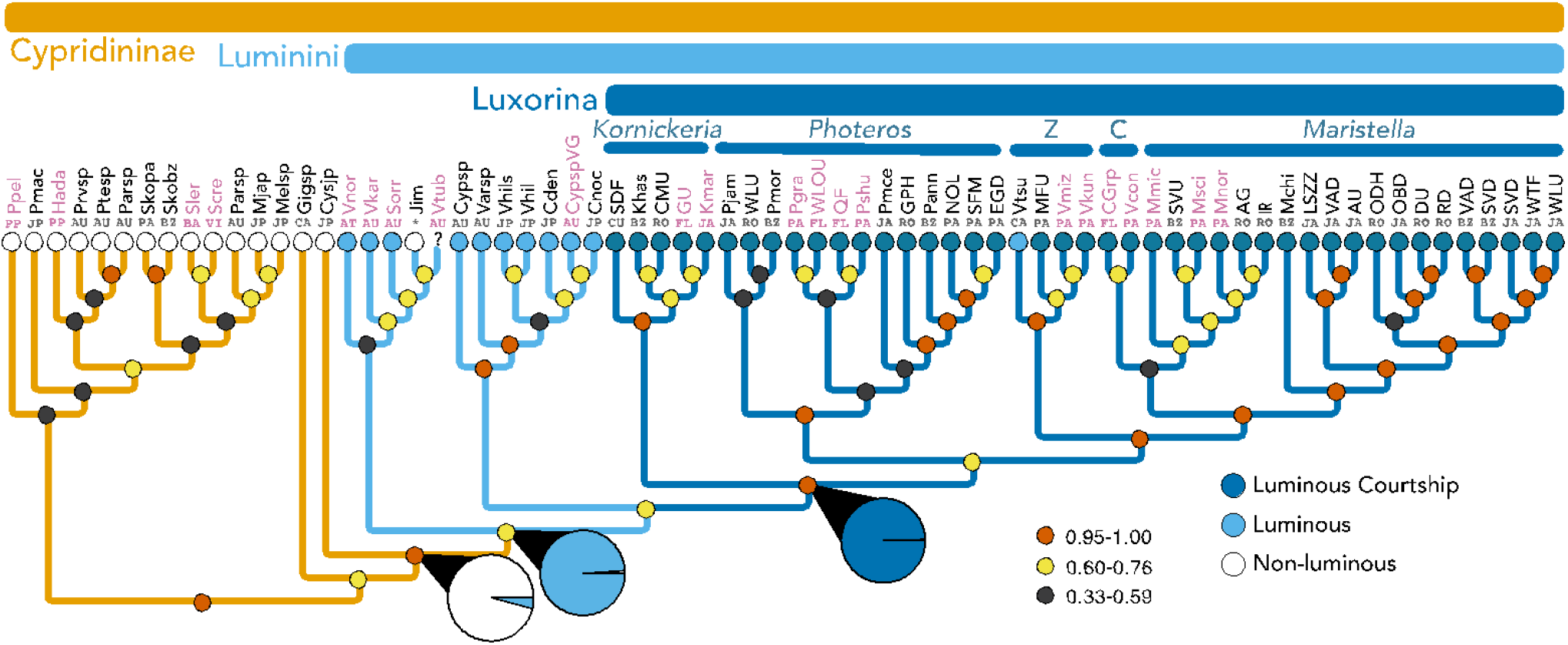
Expanded taxon sampling by adding individual mitochondrial genes, mainly from Torres and Gonzalez (2007) to the transcriptome data in Fig. 1. We inferred this species phylogeny using the multi-species-coalescent. Taxa colored pink are those that contain only mitochondrial data. Small colored circles at nodes indicate Posterior Probabilities (low=black, medium=yellow, high=red). Locality abbreviations are as in Fig. 1 and also include additional locations of VI = US Virgin Islands and FL = Florida USA. For Jimmorinia (*), we combined data from transcriptomes from a Jamaican species (*J. cf gunnari*) with mitochondrial DNA from a specimen (*Jimmorinia* sp USVI) unidentified to species from VI. Pie charts represent proportions of likelihood values for alternative ancestral states. Bioluminescence was most likely absent (white) prior to Luminini, as supported by a significantly high proportion of likelihood for absence of bioluminescence compared to presence (light blue). Significantly high likelihood proportions first appear in Luminini for bioluminescence (light blue) and Luxorina for bioluminescent courtship signaling (dark blue), compared to absence of those traits (black), which were analyzed separately as binary traits (see Supplement for full ancestral state reconstruction results).

## Discussion

How sexual selection impacts biodiversity over long evolutionary timeframes is an open question. If sexual selection affects the relative rates of extinction and speciation, it would influence the longevity of traits and clades, ultimately impacting patterns of biodiversity. However, population-based results indicate sexual selection may either increase or decrease extinction risk (Kokko and Brooks 2003; Martínez-Ruiz and Knell 2017), leading to uncertainty about the long-term persistence of sexually selected traits. Furthermore, only if consistent and conserved during evolutionary radiations should microevolutionary processes fully dictate macroevolutionary patterns. Therefore, the long-term persistence of sexually selected traits on macroevolutionary timescales is an open empirical question (Wiens and Tuschhoff 2020). Our result in Luxorina ostracods of ancient co-option of bioluminescent anti-predator signals and subsequent maintenance for courtship - lasting at least 151 Myr - provides a compelling example of a trait almost certainly under sexual selection with very long macroevolutionary persistence. The maintenance of bioluminescent courtship is counter to one line of population-level reasoning that the conspicuousness of many sexually selected traits should routinely lower fitness and increase rates of extinction and/or loss of traits under sexual selection (Promislow 1992; Tanaka 1996; Sorci et al. 1998; Doherty et al. 2003; Morrow and Fricke 2004) and instead more consistent with sexual selection reducing mutation load, increasing mean fitness, and leading to lower extinction risks (Tomkins et al. 2004; Long et al. 2012; Plesnar-Bielak et al. 2012; Lumley et al. 2015).

Besides the effects of microevolutionary processes on extinction, other factors also influence the longevity of sexually selected traits and the clades that contain them, especially biogeography and life history. The biogeography of courtship signaling in Luxorina ostracods appears restricted to the Caribbean. Therefore, Cohen and Morin (Cohen and Morin 2003) hypothesized that Luxorina is a relatively recent evolutionary radiation, yet it must be older than the formation of the Isthmus of Panama, 3.5-25 Ma (Coates et al. 1992; Farris et al. 2011; Bacon et al. 2015; O'Dea et al. 2016 and references therein). Our molecular data pushes back the origin of Luxorina and their bioluminescent courtship to about 197 (151-248) Ma (Fig. 2, Table 1), broadly consistent with the timing of initial crust formation between the Americas 154-190 Ma (Baumgartner et al. 2013). This suggests Luxorina originated and evolved on a time-scale very similar to the Proto-Caribbean + Caribbean itself, while diversifying into at least four major distinct clades (genera), each with an origin of tens of millions of years ago (Table 1), with high species diversity, and present-day distributions across most of the Caribbean (Cohen and Morin 1993, 2010; Reda et al. 2019).

The numerous islands of the Caribbean may have interacted with ecological and life history characters in Luxorina to influence species diversity and therefore longevity of the entire clade. Luxorina are shallow-water species with dispersal limited by internal fertilization, brooding, and crawl-away instars (Cohen 1983; Wakayama 2007; Gerrish and Morin 2008; Goodheart et al. 2020). Dispersal is also limited in Luxorina by micro-habitat fidelity and depth specificity (Morin 2019). Dispersal limitation and habitat specificity should increase endemic diversity in marine environments, especially in shallow waters adjacent to islands (Pinheiro et al. 2017). As one example, within the luxorine ostracod *Photeros annecohenae*, strong genetic differentiation occurs in populations <100 km apart (Reda 2019). Consistent with studies in other systems like rift lake cichlid fishes (Wagner et al. 2012), these factors point to the strong possibility of ecological opportunity interacting with sexual selection to diversify Luxorina and contribute to its longevity.

While persistence of courtship signaling since its origin over 151 Ma is the rule across dozens of Luxorina species, at least one exception does exist. *Vargula tsujii*, with no known bioluminescent courtship (Cohen and Morin 2003; Goodheart et al. 2020), is a California species now separated by the Isthmus of Panama from its closest relatives in the Caribbean, which all produce luminous courtship signals. While Morin and others have searched for luminous ostracods along the North American Pacific coast, *Vargula tsujii* remains the only known luxorine species outside the Caribbean. Most known luxorine species (~80%) in the Caribbean are collected only as adults using hand nets during brief periods of luminescent courtship, and not found baited traps or sand sweep collections. If Luxorina species in the Pacific have lost their luminescent displays or have different seasonal, lunar or diel timing for courtship behaviors, they could evade current sampling techniques. Therefore, additional Luxorina species could exist along the Pacific coast of the Americas such that extending sampling strategies could uncover additional Luxorina outside the Caribbean. For the only known luxorine from outside the Caribbean, *Vargula tsujii*, our transcriptome analyses infer a sister-group relationship to an undescribed signaling species from Panama (MFU) we found in very shallow waters near mangroves, within the provisional genus-level “Z-group”. Presumably, the *V. tsujii* lineage was originally contiguously distributed with the rest of Luxorina, until the closure of the Isthmus of Panama. *Vargula tsujii* may have lost courtship signaling after this geological event. Why *V. tsujii* lost courtship signaling is challenging to answer because it is a singular event.

Our inferences of very ancient transitions in bioluminescence traits, including long persistence of courtship signaling, depend on taxon sampling. However, better taxon sampling should not affect our main conclusion and only make these estimates older; if *Enewton* is the sister group of all other Luxorina, it would push back the origin of luminous courtship. Similarly, including species like *Vargula norvegica* and *Sheina orri* in divergence estimates adds an older node within Luminini to make the origin of bioluminescence even older. In addition, because the ttscm technique does not estimate character state changes at nodes only, it may be less affected by taxon sampling because it reconstructs distributions of ages. Therefore, the most conservative interpretation of our results is to use the 95% confidence intervals of critical nodes to make minimum estimates for character state transitions. Assuming such minimum estimates, the origin of courtship signaling is at least 151 Ma and the origin of bioluminescence is at least 197 Ma in cypridinids. Even using these very conservative minima, the origin of courtship signaling represents long persistence of a sexually selected trait and Luminini as one of the oldest origins of bioluminescence quantified thus far.

### Summary

Bioluminescence evolved in the family Cypridinidae independently from other animals, and cypridinids use a substrate, cypridinid-luciferin (Morin 2011), that is endogenous and chemically different from that of other bioluminescent organisms (Hastings 1983; Thompson et al. 1989; Kato et al. 2004, 2007). Therefore, Luminini represent a critical lineage for understanding how bioluminescence evolved (Haddock et al. 2010; Morin 2019; Goodheart et al. 2020; Lau and Oakley 2020). A phylogenetic framework including divergence time estimates provides a foundation for important questions related to molecular evolution and features important for rapid diversification, including those related to behavior (Boake et al. 2002), sexual selection (Ritchie 2007), species diversification (Ellis and Oakley 2016), and the influence of the biochemical properties of bioluminescence on evolution (Hensley et al. 2019).

The macroevolutionary durations of traits under sexual selection are difficult to predict from population-level theory, and understudied empirically. We report the maintenance of bioluminescent courtship in Luxorina ostracods for at least 151 Ma, since it was originally co-opted from bioluminescence used in anti-predator displays. Some other taxa also demonstrate long persistence of sexually selected traits (Gu et al. 2012; Bao et al. 2019; Chen and Wiens 2020; Song et al. 2020), but the macroevolutionary durations of such traits is understudied. As our understanding of the duration of sexually selected traits grows across systems, researchers will be able to address open questions about how sexual selection affects biodiversity, while disentangling other contributions such as ecological opportunity and life history characteristics.

## Acknowledgments

We acknowledge funding support from the National Science Foundation to THO (DEB-1457754, DEB-1146337), ET (DEB-1457462), GAG (DEB-1457439), JAG (BIO PRFB-1711201) and EAE (1515576 and 1702011). Permits from the Jamaican National Environment and Planning Agency (Permit Ref. no. 18/27), the Belize Fisheries Department (Permit no. 000003-16), the Honduran Department of Fish and Wildlife (Permit no. DE-MO-082-2016), the Puerto Rican Department of Natural and Environmental Resources (DRNA; Permit no. 2016-IC-113), and Panamanian Ministry of the Environment (MiAMBIENTE; Permit no. SE/A-33-17) were obtained for collections. We thank K. Osborn and J. Fergus for *Gigantocypris*. We thank Y. Mitani, Y. Ohmiya, and K. Ogoh for assistance collecting Japanese ostracods. Thanks to A. Parker for sending samples from Australia and A. Cohen for much wisdom shared through countless discussions with all of us and for confirming identification of some species.

## Data Availability Statement

Voucher samples can be accessed through the Australian Museum (AMS), National Museum of Natural History (NMNH), Santa Barbara Museum of Natural History (SBMNH), and the Museum of Comparative Zoology (MCZ) (see Table S1 for accession numbers). RNA-Seq sequence data can be accessed at the NCBI Sequence Read Archive (SRA) (see Table S1 for accession numbers). Aligned data matrices and raw tree files can be accessed at the Dryad Digital Repository (DOI: To be submitted).

