## Supplemental Figures and Tables for "Sexual signals persist over deep time: ancient co-option of bioluminescence for courtship displays in cypridinid ostracods"

#### List of Supplemental Figures

**Figure S1**. Multi-species coalescent species tree and duplication of Figure 1 for convenient comparison to additional analyses in this supplement.

**Figure S2**. Multi-species coalescent species tree (as in Fig S1) with full taxon names.

**Figure S3**. Species tree from concatenated analysis of genes from orthofinder and phylopypruner

**Figure S4**. Gene concordance values for phylogeny in S3

**Figure S5**. Bayesian relaxed molecular clock analysis and duplication of Figure 2 for convenient comparison to additional analyses in this supplement.

**Figure S6**. Bayesian relaxed molecular clock analysis as in S8 showing 95% Highest Posterior Densities at each node.

**Figure S7**. Sensitivity analysis of divergence time estimates using subsets of genes with Sortadate. We compared results from 15, 20, 25 and 30 genes, keeping all other parameters constant.

**Figure S8**. Posterior probability values of nodal support from Bayesian analysis using 15 genes selected with sortadate.

**Figure S9**. Species tree using Multi-species coalescent, using expanded taxon sampling by adding individual mitochondrial genes and duplicate of Figure 3 for convenient comparison to additional analyses in this supplement.

**Figure S10**. Species tree using Multi-species coalescent and expanded taxon sampling (as in Fig S10) with full taxon names

**Figure S11**. Species tree from concatenated analysis of only mitochondrial data alone, including 12S, 16S, and co1

**Figure S12**. Ancestral state reconstruction of presence/absence of bioluminescence in Cypridinidae.

**Figure S13**. Ancestral state reconstruction of presence/absence of bioluminescent courtship signaling in Cypridinidae.

#### List of Supplemental Tables

**Table S1**. Specimen collection and transcriptome sequencing information

**Table S2**. Length and height of carapaces to tentatively assign species to genera

**Table S3**. BUSCO scores of transcriptome assemblies

**Table S4**. Values from Gene Concordance (gCF) and Site Concordance (sCF) analyses

**Table S5**. Sample and collection data for mitochondrial data

### I. Species tree phylogenetic analyses of new transcriptomic data

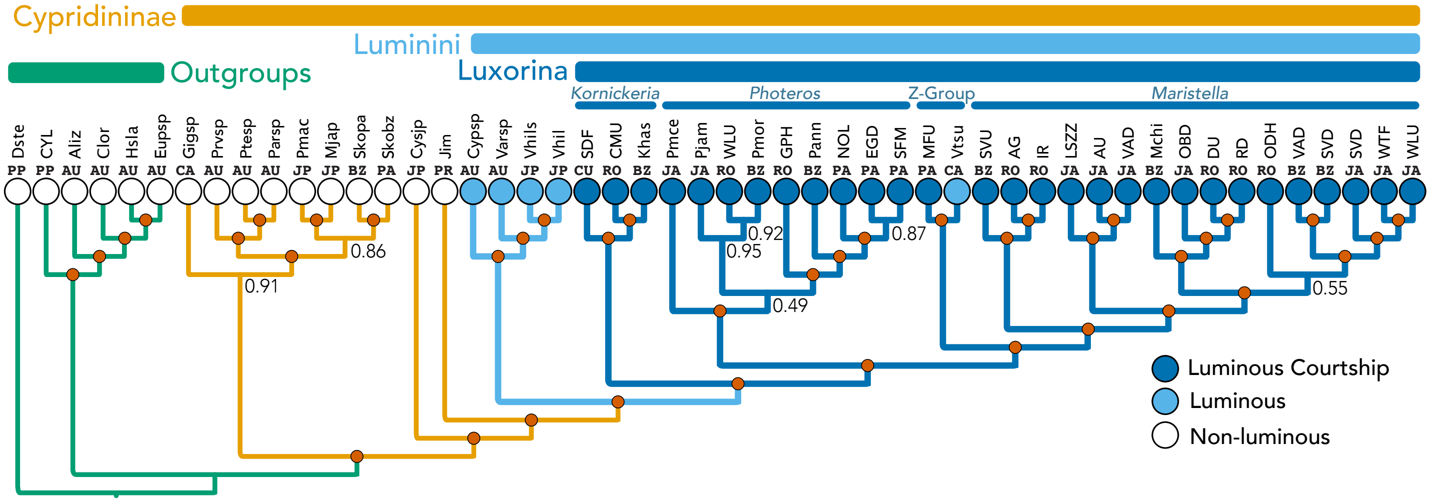


**Figure S1**. Duplication of Figure 1 for convenient comparison to additional analyses in this supplement. Species phylogeny inferred using multi-species-coalescent, employed with ASTRAL-PRO (Zhang et al. 2020) with 8485 gene trees, each of which included both orthologs and paralogs. We calculated each gene tree using fasttree 2.1.10 (Price et al. 2010) assuming a JTT+CAT model. Above the tree, we also labeled non-cypridinid Outgroups (green), and Cypridinidinae (Orange). We herein suggest a name for the bioluminescent Cypridinidae to be Tribe Luminini (after the Greek lumen for light + -ini = typical zoological tribe suffix) and a name for the clade containing signaling cypridinids to be Sub-Tribe Luxorina (lux = light + uxoriae ~= courtship + -ina = typical zoological subtribe suffix). Locality abbreviations for collecting sites of species are above each tip as follows: AU = Australia, BZ=Belize, CA=-California, USA, CU=Curacao, JA= Jamaica, JP=Japan, PA=Panama, PP=Previously Published, PR = Puerto Rico, and RO=Roatan, Honduras (see Table S1 for specific localities). Above locality names are abbreviations for species, which are expanded in Fig. S2.

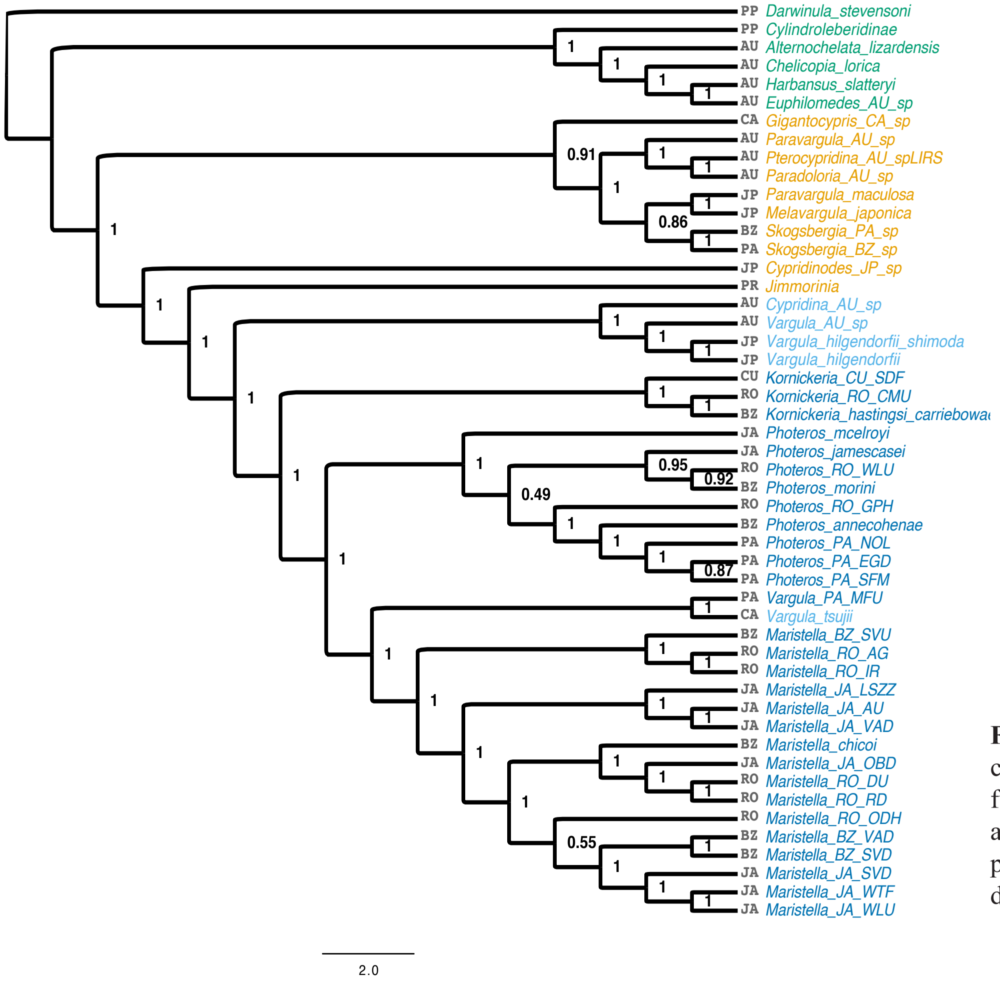


**Figure S2**. Multi-species coalescent species tree (equivalent to Figs 1, S1) with full species names rather than abbreviations. Numbers at nodes are posterior probability values. See Figs 1 and S1 for detailed methods.

**Table S1**. Specimen collection and transcriptome sequencing information

| Taxon | Country | Lat. | Long. | Voucher Museum | Voucher Accession | Transcriptome Accession | Transcriptome file name (local) | Library Prep Kit | Illumina Vers. | Sequencing Facility |
|---|---|---|---|---|---|---|---|---|---|---|
| *Alternochelata lizardensis* | Australia | -14.6689 | 145.4594 | AMS | P.92342-4 | SRX3829269 | TOCC003A | RNA TruSeq V2 | v2 PE | BGI |
| *Chelicopia lorica* | Australia | -14.6828 | 145.4481 | AMS | P.92345-8 | SRR17097280 | TOCC003B | RNA TruSeq V2 | v2 PE | BGI |
| *Cypridina sp* | Australia | | | AMS | P.92467 | SRR17097274 | TOCC004G | RNA TruSeq V2 | v2 PE | BGI |
| *Euphilomedes sp* | Australia | | | AMS | P.92379-80 | SRR17097276 | TOCC004C | RNA TruSeq V2 | v2 PE | BGI |
| *Harbansus slatteryi* | Australia | -14.6830 | 145.4495 | AMS | P.92355-61 | SRR17097279 | TOCC003F | RNA TruSeq V2 | v2 PE | BGI |
| *Paradoloria sp* | Australia | | | AMS | P.92314-5 | SRR17097273 | TOCC004H | RNA TruSeq V2 | v2 PE | BGI |
| *Paravargula sp* | Australia | | | AMS | P.92316-7 | SRR17097278 | TOCC003G | RNA TruSeq V3 | v3 PE | UC Davis Genome Center |
| *Pterocypridina spLIRS* | Australia | | | AMS | P.92319 | SRR17097271 | TOCC004I | RNA TruSeq V2 | v2 PE | BGI |
| *Vargula sp* | Australia | | | AMS | P.92349 | SRR17097275 | TOCC004F | RNA TruSeq V2 | v2 PE | BGI |
| *Kornickeria hastingsi carrie-bowae* | Belize | 16.8116 | -88.0824 | NMNH | No voucher | SRR10860879 | Korn_hast_c | Prepared at Novogene | Novo-gene | Novogene |
| *Kornickeria hastingsi carrie-bowae* | Belize | 16.8116 | -88.0824 | NMNH | No voucher | SRR10860878 | Kornickeria_hastingsii_carrie-bowae_Rep2 | RNA TruSeq V2 | v2 PE | Functional Genomics Lab |
| *Maristella BZ SVU* | Belize | 16.8116 | -88.0824 | NMNH | USNM 1659658 | SRR10811643 | Sample_TOCC001C_index3 | RNA TruSeq V2 | v3 PE | Functional Genomics Lab |
| *Maristella BZ SVU* | Belize | 16.8116 | -88.0824 | NMNH | USNM 1659658 | SRR10811644 | B_SVU | NEBNext® Ultra™ RNA | v3 PE | Novogene |
| *Maristella BZ SVD* | Belize | 16.8116 | -88.0824 | NMNH | USNM 1659659 | SRR10860875 | Maristella_SVD | NEBNext® Ultra™ RNA | v3 PE | Novogene |
| *Maristella BZ SVD* | Belize | 16.8116 | -88.0824 | NMNH | USNM 1659659 | SRR10860874 | Maristella_SVD_Rep2 | RNA TruSeq v2 | v2 PE | BGI |
| *Maristella BZ VAD* | Belize | 16.8065 | -88.0790 | NMNH | USNM 1659657 | SRR17097283 | BZ-VAD2_S4 | RNA TruSeq V3 | v3 PE | UC Davis Genome Center |
| *Maristella BZ VAD* | Belize | 16.8065 | -88.0790 | NMNH | USNM 1659657 | SRR17097272 | BZ-VAD_S4 | RNA TruSeq V3 | v3 PE | UC Davis Genome Center |
| *Maristella BZ VAD* | Belize | 16.8065 | -88.0790 | NMNH | USNM 1659657 | SRR17097270 | BZVAD | RNA TruSeq V3 | v3 PE | UC Davis Genome Center |
| *Maristella chicoi* | Belize | 16.8116 | -88.0824 | NMNH | No voucher | SRR10811640 | B_MSH | NEBNext® Ultra™ RNA | v3 PE | Novogene |
| *Maristella chicoi* | Belize | 16.8116 | -88.0824 | NMNH | No voucher | SRR10811639 | BzMSH | RNA TruSeq V3 | v3 PE | Functional Genomics Lab |
| *Photeros annecohenae* | Belize | 16.8116 | -88.0824 | NMNH | No voucher | SRR10860880 | Gruber_Pannecohenae | RNA TruSeq v3 | v3 PE | New York Genome Center |
| *Photeros morini* | Belize | 16.8116 | -88.0824 | NMNH | No voucher | SRR10860877 | B_Pmorin | NEBNext® Ultra™ RNA | v3 PE | Novogene |
| *Photeros morini* | Belize | 16.8116 | -88.0824 | NMNH | No voucher | SRR10860876 | Sample_TOCC001B_index2 | RNA TruSeq V2 | v3 PE | Functional Genomics Lab |
| *Skogsbergia sp BZG* | Belize | 16.7470 | -87.8097 | No voucher | No voucher | SRR17097268 | GlovSkogs | RNA TruSeq V2 | v3 PE | BGI |
| *Kornickeria CU SDF* | Curacao | 12.1094 | -68.9545 | SBMNH | SBMNH 658185 | SRR17097269 | CURA | Prepared at Novogene | Novo-gene | Novogene |
| *Kornickeria RO CMU* | Honduras | 16.3581 | -86.4329 | NMNH | USNM 1659663 | SRR17097291 | R-CMU_S63_L008 | RNA TruSeq V3 | v3 PE | UC Davis Genome Center |

| | | | | | | | | | | |
|---|---|---|---|---|---|---|---|---|---|---|
| *Kornickeria RO CMU* | Honduras | 16.3581 | -86.4329 | NMNH | USNM 1659663 | SRR17097292 | R-CMU_S5 | RNA TruSeq V3 | v3 PE | UC Davis Genome Center |
| *Maristella RO AG* | Honduras | 16.8065 | -88.0790 | NMNH | USNM 1659662 | SRR10811636 | EETO030418-RO_AG | NEBNext® Ultra™ RNA | v3 PE | UCSB-BNL |
| *Maristella RO DU* | Honduras | 16.8065 | -88.0790 | NMNH | USNM 1659667 | SRR17097289 | R-DU_S60 | RNA TruSeq V3 | v3 PE | UC Davis Genome Center |
| *Maristella RO DU* | Honduras | 16.8065 | -88.0790 | NMNH | USNM 1659667 | SRR17097290 | R-DU_S3 | RNA TruSeq V3 | v3 PE | UC Davis Genome Center |
| *Maristella RO DU* | Honduras | 16.8065 | -88.0790 | NMNH | USNM 1659667 | SRR17097284 | R2016DU | RNA TruSeq V3 | v3 PE | UC Davis Genome Center |
| *Maristella RO IR* | Honduras | 16.8065 | -88.0790 | NMNH | USNM 1659665 | SRR12672004 | R-IR_S66_L008 | RNA TruSeq V3 | v3 PE | UC Davis Genome Center |
| *Maristella RO ODH* | Honduras | 16.8065 | -88.0790 | NMNH | USNM 1659661 | SRR12672003 | R_ODH | NEBNext® Ultra™ RNA | v3 PE | Novogene |
| *Maristella RO ODH* | Honduras | 16.8065 | -88.0790 | NMNH | USNM 1659661 | SRR17097288 | R-OHD_S65_L008 | RNA TruSeq V3 | v3 PE | UC Davis Genome Center |
| *Maristella RO RD* | Honduras | 16.3581 | -86.4329 | NMNH | USNM 1659660 | SRR17097285 | R-RD2_S1 | RNA TruSeq V3 | v3 PE | UC Davis Genome Center |
| *Maristella RO RD* | Honduras | 16.3581 | -86.4329 | NMNH | USNM 1659660 | SRR17097287 | R-RD_S1 | RNA TruSeq V3 | v3 PE | UC Davis Genome Center |
| *Maristella RO RD* | Honduras | 16.3581 | -86.4329 | NMNH | USNM 1659660 | SRR17097287 | R-RD_S59 | RNA TruSeq V3 | v3 PE | UC Davis Genome Center |
| *Photeros RO GPH* | Honduras | 16.4027 | -86.4090 | NMNH | USNM 1659666 | SRR17097293 | R_GPH | NEBNext® Ultra™ RNA | v3 PE | UCSB-BNL |
| *Photeros RO GPH* | Honduras | 16.4027 | -86.4090 | NMNH | USNM 1659666 | SRR17097282 | RO2015GPH | RNA TruSeq V3 | v3 PE | UC Davis Genome Center |
| *Photeros RO WLU* | Honduras | 16.8065 | -88.0790 | NMNH | USNM 1659664 | SRR10811635 | EETO030418-RO_WLU | NEBNext® Ultra™ RNA | v3 PE | UCSB-BNL |
| *Jimmorinia gunnari* | Jamaica | 18.4700 | -77.4100 | NMNH | No voucher | SRR17097315 | J_JimGun-43942958 | NEBNext® Ultra™ RNA | v3 PE | UCSB-BNL |
| *Jimmorinia gunnari* | Jamaica | 18.4700 | -77.4100 | NMNH | No voucher | SRR17097306 | JAM_JIMMOR | NEBNext® Ultra™ RNA | v3 PE | Novogene |
| *Maristella JA AU* | Jamaica | 18.4723 | -77.4095 | NMNH | USNM 1659654 | SRR17097312 | J-AU_S56_L007 | RNA TruSeq V3 | v3 PE | UC Davis Genome Center |
| *Maristella JA LSZZ* | Jamaica | 18.4715 | -77.4148 | NMNH | USNM 1659653 | SRR17097300 | LSZZ_S3 | RNA TruSeq V3 | v3 PE | UC Davis Genome Center |
| *Maristella JA LSZZ* | Jamaica | 18.4715 | -77.4148 | NMNH | USNM 1659653 | SRR17097304 | JAM_LSZZ | RNA TruSeq V3 | v3 PE | UC Davis Genome Center |
| *Maristella JA OBD* | Jamaica | 18.4723 | -77.4095 | NMNH | No voucher | SRR17097310 | J-OBD_S52 | RNA TruSeq V3 | v3 PE | UC Davis Genome Center |
| *Maristella JA OBD* | Jamaica | 18.4723 | -77.4095 | NMNH | No voucher | SRR17097314 | J_OBD | NEBNext® Ultra™ RNA | v3 PE | Novogene |
| *Maristella JA SVD* | Jamaica | 18.4802 | -77.4460 | NMNH | USNM 1659651 | SRR17097313 | J_SVD | NEBNext® Ultra™ RNA | v3 PE | Novogene |
| *Maristella JA SVD* | Jamaica | 18.4802 | -77.4460 | NMNH | USNM 1659651 | SRR17097309 | J-SVD_S53 | RNA TruSeq V3 | v3 PE | UC Davis Genome Center |
| *Maristella JA VAD* | Jamaica | 18.4737 | -77.4169 | NMNH | USNM 1659652 | SRR10811642 | J-VAD_S55 | RNA TruSeq V3 | v3 PE | UC Davis Genome Center |
| *Maristella JA VAD* | Jamaica | 18.4737 | -77.4169 | NMNH | USNM 1659652 | SRR10811641 | J_VAD-43931095 | NEBNext® Ultra™ RNA | v3 PE | UCSB-BNL |

| | | | | | | | | | | |
|---|---|---|---|---|---|---|---|---|---|---|
| *Maristella JA WLU* | Jamaica | 18.4723 | -77.4095 | NMNH | USNM 1659647 | SRR17097308 | J-WLU_S67_L008 | RNA TruSeq V3 | v3 PE | UC Davis Genome Center |
| *Maristella JA WTF* | Jamaica | 18.4733 | -77.4128 | NMNH | USNM 1659655 | SRR17097307 | J-WTF_S54_L007 | RNA TruSeq V3 | v3 PE | UC Davis Genome Center |
| *Photeros jamescasei* | Jamaica | 18.4737 | -77.4169 | NMNH | USNM 1659648 | SRR17097311 | J-Jamescasei2_S5 | RNA TruSeq V3 | v3 PE | UC Davis Genome Center |
| *Photeros jamescasei* | Jamaica | 18.4737 | -77.4169 | NMNH | USNM 1659648 | SRR17097267 | J_Jamescasei-43938115 | NEBNext® Ultra™ RNA | v3 PE | UCSB-BNL |
| *Photeros mcelroyi* | Jamaica | 18.4751 | -77.4055 | NMNH | USNM 1659650 | SRR10811638 | J-MacElroyi_S2 | RNA TruSeq V3 | v3 PE | UC Davis Genome Center |
| *Photeros mcelroyi* | Jamaica | 18.4751 | -77.4055 | NMNH | USNM 1659650 | SRR10811637 | J-MacElroyi2_S2 | RNA TruSeq V3 | v3 PE | UC Davis Genome Center |
| *Photeros mcelroyi* | Jamaica | 18.4751 | -77.4055 | NMNH | USNM 1659650 | SRR10811645 | JMACELROY | RNA TruSeq V3 | v3 PE | UC Davis Genome Center |
| *Cypridinodes sp* | Japan | 34.6737 | 138.9743 | SBMNH | SBMNH 658174 | - | jpcyp2b | NEBNext® Ultra™ RNA | v3 PE | Novogene |
| *Melavargula japonica* | Japan | 34.6737 | 138.9743 | SBMNH | SBMNH 658182 | SRR17097303 | Jp_Melavar22 | NEBNext® Ultra™ RNA | v3 PE | Novogene |
| *Paravargula maculosa* | Japan | 34.6737 | 138.9743 | SBMNH | SBMNH 658180 | SRR17097305 | 3_S8_L001 | RNA TruSeq V3 | v3 PE | UC Davis Genome Center |
| *Paravargula maculosa* | Japan | 34.6737 | 138.9743 | SBMNH | SBMNH 658180 | SRR17097294 | 3-2_S9_L001 | RNA TruSeq V3 | v3 PE | UC Davis Genome Center |
| *Vargula hilgendorfii* | Japan | | | No voucher | No voucher | SRR17097277 | TOCC003H | RNA TruSeq V2 | v2 PE | BGI |
| *Vargula hilgendorfii 2* | Japan | | | No voucher | No voucher | SRR17097317 | 1_S6 | RNA TruSeq V3 | v3 PE | UC Davis Genome Center |
| *Vargula hilgendorfii 2* | Japan | | | No voucher | No voucher | SRR17097316 | 1-2_S7 | RNA TruSeq V3 | v3 PE | UC Davis Genome Center |
| *PA MFU* | Panama | 9.3313 | -82.2533 | NHNM | No voucher | SRR17097296 | PMMFU | Prepared at Novogene | Novo-gene | Novogene |
| *Photeros PA EGD* | Panama | 9.3326 | -82.2539 | SBMNH | SBMNH 658179 | SRR17097299 | Panama_Photeros_EGD__EGD_Niko | Prepared at Novogene | v3 PE | Novogene |
| *Photeros PA NOL* | Panama | 9.3546 | -82.2587 | SBMNH | SBMNH 658177 | SRR17097298 | Panama_Photeros_NOL__NOL_Niko | Prepared at Novogene | v3 PE | Novogene |
| *Photeros PA SFM* | Panama | 9.3327 | -82.2536 | SBMNH | SBMNH 658201 | SRR17097297 | Panama_Photeros_SFM__SFM_Niko | Prepared at Novogene | v3 PE | Novogene |
| *Skogsbergia sp PA* | Panama | 9.3300 | -82.2500 | | No voucher | SRR17097295 | PSKOG | Prepared at Novogene | Novo-gene | Novogene |
| *Gigantocypris sp* | USA | | | | No voucher | SRR17097301 | L4_Gigantocypris_MBARI | RNA TruSeq V2 | v3 PE | BGI |
| *Vargula tsujii* | USA | 33.4451 | -118.4845 | SBMNH | SBMNH 660969 | SRR1269674 | TOCC003I | RNA TruSeq V2 | v2 PE | BGI |
| *Cylindroleberidinae* | Belize | 16.8030 | -88.0810 | MCZ | MCZ IZ 74242 | SRR4113495 | | | | |
| *Darwinula stevensoni* | Belgium | Unknown | | Unknown | Unknown | ENA PRJEB38362 | | | | |

**Table S2**. Carapace measurements

| Species Designation | Collector | Meas. by | Lat. | Long. | N (len/ eye/keel) | Len (mm) | Min. Len | Max. Len | Height (mm) | Min. Height | Max. Height | Eye (mm) | Min. Eye | Max. Eye | Keel (mm) | Min. Keel | Max. Keel | L/H |
|---|---|---|---|---|---|---|---|---|---|---|---|---|---|---|---|---|---|---|
| *Kornickeria CU SDF* | JAG/THO | | 12.1094 | -68.9545 | | | | | | | | | | | | | | |
| *Kornickeria hastingsi carriebowie* | GAG/JGM | GAG | 16.8116 | -88.0824 | 37/15/15 | 1.80 | 1.72 | 1.86 | 1.01 | 0.96 | 1.06 | 0.25 | 0.22 | 0.28 | 0.12 | 0.10 | 0.12 | 1.774 |
| *Kornickeria RO CMU* | GAG/EAE | GAG | 16.3581 | -86.4329 | 67/28/28 | 1.77 | 1.72 | 1.86 | 1.00 | 0.96 | 1.05 | 0.20 | 0.16 | 0.25 | 0.12 | 0.08 | 0.18 | 1.775 |
| *Maristella BZ SVD* | GAG/JGM | GAG | 16.8116 | -88.0824 | 19/13/13 | 1.76 | 1.68 | 1.86 | 1.06 | 0.98 | 1.10 | 0.18 | 0.16 | 0.20 | 0.07 | 0.06 | 0.10 | 1.666 |
| *Maristella BZ SVU* | GAG/JGM | GAG | 16.8116 | -88.0824 | 71/15/15 | 2.17 | 2.06 | 2.28 | 1.31 | 1.22 | 1.42 | 0.27 | 0.24 | 0.36 | 0.17 | 0.14 | 0.22 | 1.664 |
| *Maristella BZ VAD* | GAG/EAE | GAG | 16.8065 | -88.0790 | 35/20/20 | 1.87 | 1.78 | 1.93 | 1.11 | 1.05 | 1.15 | 0.23 | 0.20 | 0.28 | 0.11 | 0.10 | 0.13 | 1.690 |
| *Maristella chicoi* | GAG/JGM | GAG | 16.8116 | -88.0824 | 68/32/32 | 1.62 | 1.56 | 1.70 | 0.96 | 0.92 | 1.00 | 0.22 | 0.16 | 0.26 | 0.07 | 0.04 | 0.08 | 1.686 |
| *Maristella JA AU* | GAG/JPF | GAG | 18.4723 | -77.4095 | 34/34/34 | 2.17 | 2.10 | 2.23 | 1.23 | 1.15 | 1.28 | 0.25 | 0.20 | 0.28 | 0.12 | 0.10 | 0.15 | 1.772 |
| *Maristella JA LSZZ* | NMH/JGM/ TJR | GAG | 18.4715 | -77.4148 | 18/18/18 | 1.70 | 1.63 | 1.75 | 1.03 | 1.00 | 1.05 | 0.19 | 0.18 | 0.20 | 0.10 | 0.08 | 0.13 | 1.650 |
| *Maristella JA OBD* | GAG/TJR/ JGM | GAG | 18.4723 | -77.4095 | 42/27/27 | 1.79 | 1.75 | 1.85 | 1.06 | 1.00 | 1.10 | 0.21 | 0.20 | 0.25 | 0.08 | 0.08 | 0.13 | 1.692 |
| *Maristella JA SVD* | GAG/JPF/ NMH | GAG | 18.4802 | -77.4460 | 59/49/49 | 1.95 | 1.88 | 2.03 | 1.15 | 1.10 | 1.20 | 0.22 | 0.20 | 0.25 | 0.09 | 0.08 | 0.10 | 1.700 |
| *Maristella JA VAD* | TJR/JPF/ JGM | GAG | 18.4737 | -77.4169 | 47/28/28 | 2.18 | 2.08 | 2.25 | 1.21 | 1.15 | 1.25 | 0.23 | 0.18 | 0.25 | 0.11 | 0.10 | 0.15 | 1.807 |
| *Maristella JA WLU* | TJR/GAG/ NMH/JPF | GAG | 18.4723 | -77.4095 | 55/26/26 | 1.92 | 1.88 | 1.98 | 1.13 | 1.10 | 1.20 | 0.22 | 0.20 | 0.25 | 0.11 | 0.10 | 0.13 | 1.694 |
| *Maristella JA WTF* | GAG | GAG | 18.4733 | -77.4128 | 12/12/12 | 1.89 | 1.83 | 1.95 | 1.13 | 1.08 | 1.18 | 0.22 | 0.20 | 0.23 | 0.10 | 0.10 | 0.13 | 1.670 |
| *Maristella RO AG* | GAG/EAE | GAG | 16.8065 | -88.0790 | 31/31/31 | 2.29 | 2.16 | 2.40 | 1.39 | 1.30 | 1.43 | 0.22 | 0.20 | 0.25 | 0.17 | 0.13 | 0.20 | 1.649 |
| *Maristella RO DU* | GAG/JGM/ EAE | GAG | 16.8065 | -88.0790 | 54/34/32 | 1.63 | 1.58 | 1.73 | 0.98 | 0.94 | 1.05 | 0.21 | 0.18 | 0.28 | 0.09 | 0.06 | 0.10 | 1.663 |
| *Maristella RO IR* | GAG/NMH | GAG | 16.8065 | -88.0790 | 52/30/30 | 2.28 | 2.22 | 2.38 | 1.39 | 1.35 | 1.44 | 0.23 | 0.20 | 0.25 | 0.15 | 0.12 | 0.18 | 1.646 |
| *Maristella RO ODH* | GAG/JGM | GAG | 16.8065 | -88.0790 | 40/20/20 | 1.69 | 1.64 | 1.76 | 0.99 | 0.96 | 1.04 | 0.20 | 0.16 | 0.22 | 0.08 | 0.06 | 0.10 | 1.711 |
| *Maristella RO RD* | JGM/NMH | GAG | 16.3581 | -86.4329 | 51/31/31 | 1.64 | 1.58 | 1.70 | 0.98 | 0.94 | 1.03 | 0.18 | 0.14 | 0.23 | 0.08 | 0.06 | 0.10 | 1.670 |
| *PA MFU* | JGM | GAG | 9.3313 | -82.2533 | 30/20/20 | 1.66 | 1.58 | 1.70 | 1.00 | 0.95 | 1.05 | 0.19 | 0.18 | 0.20 | 0.08 | 0.08 | 0.10 | 1.654 |
| *Photeros annecohenae* | GAG/JGM | JGM | 16.8116 | -88.0824 | 121/15/15 | 1.62 | 1.50 | 1.70 | 1.02 | 0.96 | 1.08 | 0.22 | 0.18 | 0.26 | 0.12 | 0.08 | 0.16 | 1.591 |
| *Photeros jamescasei* | JPF/GAG/ JGM | GAG | 18.4737 | -77.4169 | 36/30/30 | 2.07 | 1.98 | 2.13 | 1.31 | 1.23 | 1.35 | 0.27 | 0.25 | 0.30 | 0.14 | 0.10 | 0.15 | 1.576 |
| *Photeros mcelroyi* | NMH/GAG/ JGM | GAG | 18.4751 | -77.4055 | 41/35/35 | 1.92 | 1.85 | 2.00 | 1.22 | 1.18 | 1.25 | 0.27 | 0.23 | 0.30 | 0.13 | 0.10 | 0.18 | 1.576 |
| *Photeros morini* | GAG/JGM | GAG | 16.8116 | -88.0824 | 45/15/15 | 2.06 | 1.94 | 2.14 | 1.28 | 1.20 | 1.36 | 0.37 | 0.34 | 0.40 | 0.11 | 0.08 | 0.12 | 1.604 |
| *Photeros PA EGD* | NMH/JGM | GAG | 9.3326 | -82.2539 | 18/16/16 | 1.63 | 1.58 | 1.65 | 1.04 | 1.03 | 1.08 | 0.18 | 0.18 | 0.20 | 0.10 | 0.08 | 0.13 | 1.557 |
| *Photeros PA NOL* | THO, NMH | THO | 9.3546 | -82.2587 | 14/14/14 | 1.63 | 1.55 | 1.67 | 1.04 | 1.01 | 1.07 | 0.21 | 0.19 | 0.25 | 0.10 | 0.08 | 0.13 | 1.567 |
| *Photeros PA SFM* | NMH/JGM | GAG/ EAE | 9.3327 | -82.2536 | 60/50/50 | 1.70 | 1.63 | 1.78 | 1.08 | 1.03 | 1.13 | 0.19 | 0.16 | 0.23 | 0.09 | 0.05 | 0.12 | 1.581 |
| *Photeros RO GPH* | GAG | GAG | 16.4027 | -86.4090 | 40/20/20 | 1.68 | 1.60 | 1.74 | 1.04 | 1.00 | 1.08 | 0.23 | 0.22 | 0.26 | 0.09 | 0.08 | 0.10 | 1.621 |
| *Photeros RO WLU* | GAG/JGM/ NMH | GAG | 16.8065 | -88.0790 | 53/31/31 | 1.98 | 1.82 | 2.17 | 1.22 | 1.10 | 1.43 | 0.24 | 0.20 | 0.30 | 0.13 | 0.10 | 0.17 | 1.619 |

**Table S3**. Metrics of transcriptome completeness

| Species | BUSCO v3 Complete (Max 1066 arth_db9) | Number of Contigs | N50 | Transdecoder.v3 Complete ORFs |
|---|---|---|---|---|
| *Alternochelata lizardensis* | 316 | 8573 | 367 | 1443 |
| *Chelicopia lorica* | 436 | 111956 | 506 | 2736 |
| *Cylindroleberidinae* | 755 | 833810 | 389 | 15389 |
| *Cypridina sp* | 758 | 232667 | 595 | 6158 |
| *Cypridinodes sp* | 622 | 220069 | 430 | 3186 |
| *Darwinula stevensoni* | 928 | 62117 | 56422 | 14612 |
| *Euphilomedes sp* | 559 | 67419 | 444 | 5676 |
| *Gigantocypris sp* | 458 | 60294 | 545 | 3050 |
| *Harbansus slatteryi* | 623 | 112493 | 649 | 3958 |
| *Jimmorinia gunnari* | 47 | 25397 | 284 | 145 |
| *Kornickeria CU SDF* | 525 | 123953 | 391 | 2877 |
| *Kornickeria hastingsi carriebowae* | 675 | 131518 | 520 | 5040 |
| *Kornickeria RO CMU* | 914 | 459761 | 441 | 12132 |
| *Maristella BZ SVD* | 465 | 97454 | 540 | 3541 |
| *Maristella BZ SVU* | 318 | 96641 | 408 | 1507 |
| *Maristella BZ VAD* | 573 | 353073 | 368 | 4996 |
| *Maristella chicoi* | 344 | 166512 | 406 | 23425 |
| *Maristella JA AU* | 781 | 223647 | 522 | 7777 |
| *Maristella JA LSZZ* | 355 | 308104 | 326 | 2534 |
| *Maristella JA OBD* | 738 | 164684 | 580 | 8174 |
| *Maristella JA SVD* | 764 | 225704 | 516 | 7966 |
| *Maristella JA VAD* | 675 | 307747 | 386 | 6187 |
| *Maristella JA WLU* | 896 | 273365 | 608 | 12250 |
| *Maristella JA WTF* | 712 | 172775 | 561 | 7629 |
| *Maristella RO AG* | 861 | 578384 | 396 | 10812 |
| *Maristella RO DU* | 871 | 326035 | 430 | 7465 |
| *Maristella RO IR* | 584 | 169966 | 469 | 4860 |
| *Maristella RO ODH* | 803 | 202233 | 577 | 8765 |
| *Maristella RO RD* | 836 | 305801 | 486 | 8995 |
| *Melavargula japonica* | 96 | 66028 | 314 | 305 |
| *PA MFU* | 454 | 110867 | 400 | 3022 |
| *Paradoloria sp* | 755 | 168668 | 649 | 6668 |
| *Paravargula maculosa* | 251 | 52293 | 431 | 1344 |
| *Paravargula sp* | 540 | 124555 | 650 | 3086 |
| *Photeros annecohenae* | 843 | 330459 | 647 | 13707 |
| *Photeros jamescasei* | 6 | 5044 | 254 | 32 |
| *Photeros mcelroyi* | 148 | 197717 | 304 | 935 |
| *Photeros morini* | 197 | 7576 | 262 | 985 |
| *Photeros PA EGD* | 662 | 178943 | 697 | 11910 |
| *Photeros PA NOL* | 710 | 195227 | 683 | 9482 |
| *Photeros PA SFM* | 746 | 188191 | 742 | 11538 |
| *Photeros RO GPH* | 455 | 385152 | 365 | 4255 |
| *Photeros RO WLU* | 860 | 395655 | 462 | 10779 |
| *Pterocypridina spLIRS* | 514 | 145874 | 576 | 3926 |
| *Skogsbergia sp BZG* | 122 | 59380 | 365 | 8101 |
| *Skogsbergia sp PA* | 676 | 152911 | 402 | 3832 |
| *Vargula hilgendorfii* | 336 | 51891 | 561 | 1184 |
| *Vargula hilgendorfii sp 2* | 292 | 45009 | 400 | 7576 |
| *Vargula sp* | 268 | 51872 | 488 | 970 |
| *Vargula tsujii* | 607 | 122949 | 573 | 4317 |

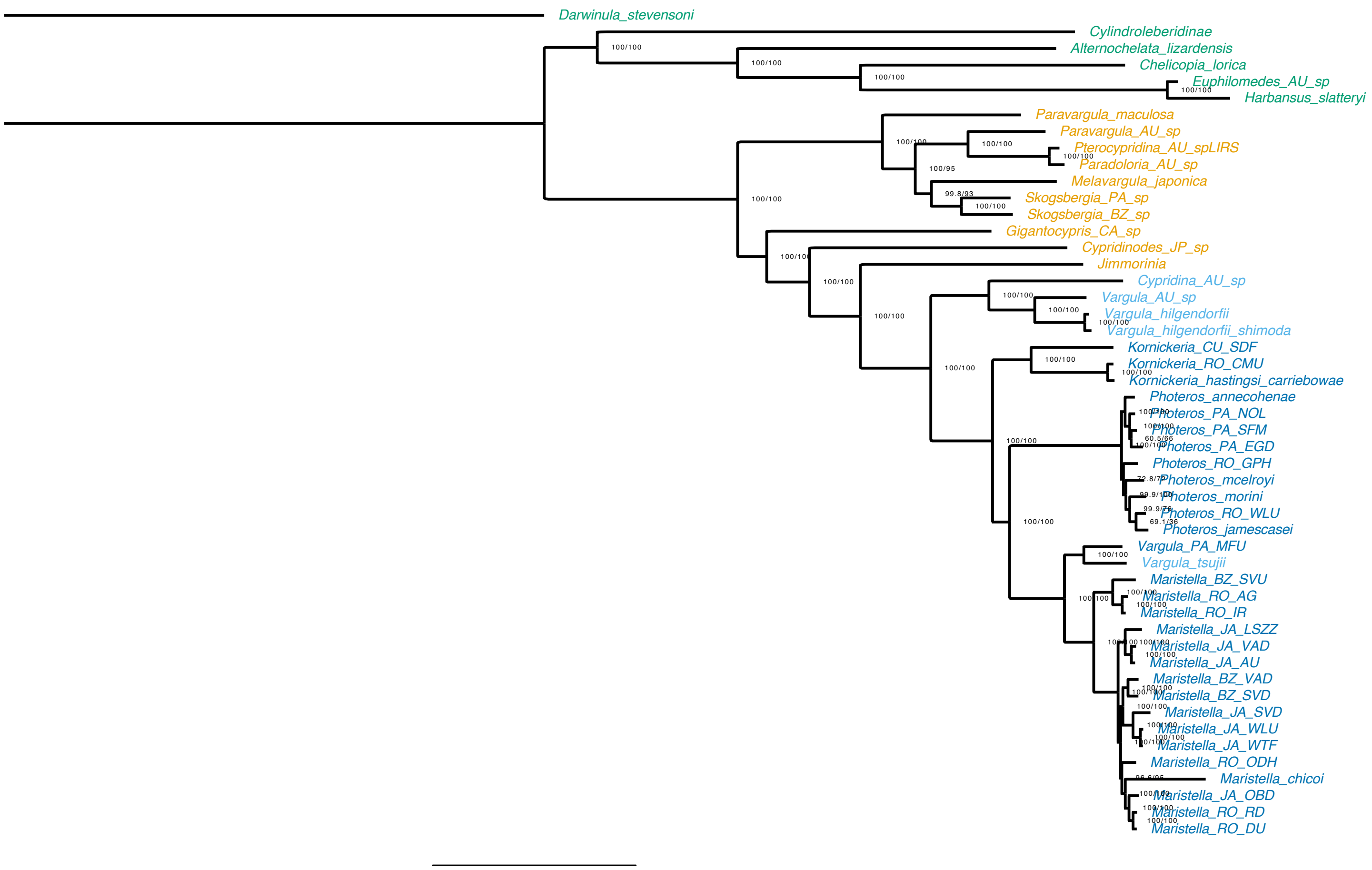


**Figure S3**. Concatenated analyses of 2675 orthologs determined with Orthofinder 2.5.2 and Phylopypruner 0.9.7 using Maximum Likelihood implemented in IQ-TREE v2.0.3, with additional options for -bnni and -alrt (set to 1000) for 1000 bootstrap replicates.

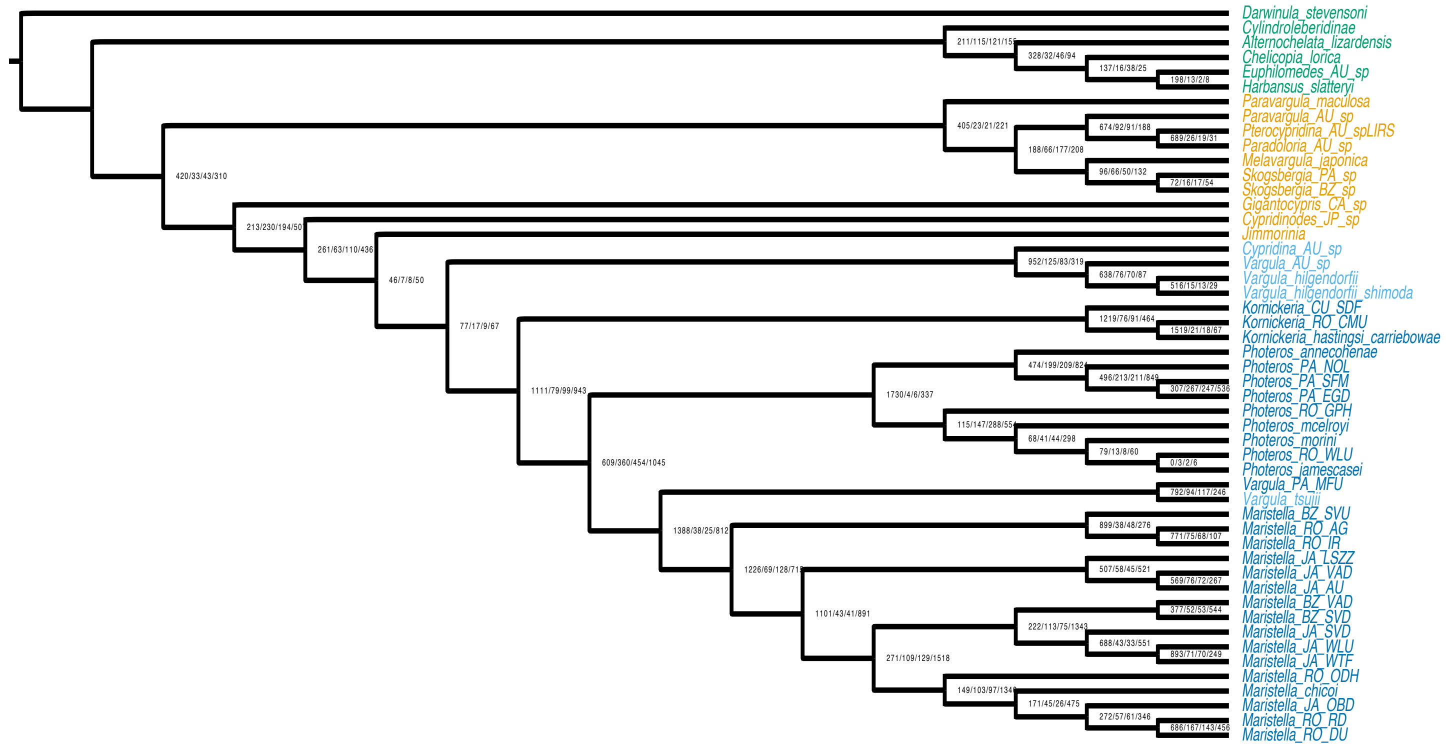


**Figure S4** - Gene concordance analysis of 2675 gene families containing only orthologs, determined with Orthofinder 2.5.2 and Phylopypruner 0.9.7, implemented in IQ-TREE v2.0.3 with an additional option set (--scf 100).

Table S4. Values from Gene Concordance (gCF) and Site Concordance (sCF) analyses implemented in IQ-TREE using orthofinder/phylopypruner orthologs.

| ID | gCF | gCF_N | gDF1 | gDF1_N | gDF2 | gDF2_N | gDFP | gDFP_N | gN | sCF | sCF_N | sDF1 | sDF1_N | sDF2 | sDF2_N | sN | Bootstrap |
|---|---|---|---|---|---|---|---|---|---|---|---|---|---|---|---|---|---|
| **51** | 47.25 | 686 | 11.5 | 167 | 9.85 | 143 | 31.4 | 456 | 1452 | 68.1 | 437.6 | 18.06 | 96.15 | 13.84 | 89.84 | 623.59 | 100/100 |
| **52** | 36.96 | 272 | 7.74 | 57 | 8.29 | 61 | 47.01 | 346 | 736 | 80.12 | 633.09 | 11.01 | 76.07 | 8.87 | 59.43 | 768.59 | 100/100 |
| **53** | 23.85 | 171 | 6.28 | 45 | 3.63 | 26 | 66.25 | 475 | 717 | 28.37 | 206.13 | 60.17 | 466.27 | 11.47 | 81.54 | 753.94 | 100/100 |
| **54** | 8.82 | 149 | 6.1 | 103 | 5.74 | 97 | 79.34 | 1340 | 1689 | 30.24 | 219.95 | 41.2 | 305 | 28.56 | 206.12 | 731.07 | 96.6/95 |
| **55** | 12.66 | 222 | 6.45 | 113 | 4.28 | 75 | 76.61 | 1343 | 1753 | 45.1 | 298.65 | 28.61 | 167.75 | 26.29 | 157.63 | 624.03 | 100/100 |
| **56** | 52.32 | 688 | 3.27 | 43 | 2.51 | 33 | 41.9 | 551 | 1315 | 67.39 | 509.67 | 16.81 | 92.63 | 15.79 | 92.88 | 695.18 | 100/100 |
| **57** | 69.6 | 893 | 5.53 | 71 | 5.46 | 70 | 19.41 | 249 | 1283 | 87.75 | 778.13 | 4.15 | 31.1 | 8.11 | 55.96 | 865.19 | 100/100 |
| **58** | 36.74 | 377 | 5.07 | 52 | 5.17 | 53 | 53.02 | 544 | 1026 | 60.84 | 418.38 | 20.13 | 102.94 | 19.03 | 116.47 | 637.79 | 100/100 |
| **59** | 13.37 | 271 | 5.38 | 109 | 6.36 | 129 | 74.89 | 1518 | 2027 | 33.92 | 234.39 | 27.83 | 178.07 | 38.25 | 260.09 | 672.55 | 100/100 |
| **60** | 53.03 | 1101 | 2.07 | 43 | 1.97 | 41 | 42.92 | 891 | 2076 | 70.35 | 826.21 | 18.86 | 162.84 | 10.79 | 114.49 | 1103.54 | 100/100 |
| **61** | 71.29 | 899 | 3.01 | 38 | 3.81 | 48 | 21.89 | 276 | 1261 | 75.96 | 782.18 | 13.35 | 105.62 | 10.69 | 102.14 | 989.94 | 100/100 |
| **62** | 75.51 | 771 | 7.35 | 75 | 6.66 | 68 | 10.48 | 107 | 1021 | 83.42 | 733.13 | 10.67 | 72.81 | 5.91 | 45.26 | 851.2 | 100/100 |
| **63M** | 57.34 | 1226 | 3.23 | 69 | 5.99 | 128 | 33.44 | 715 | 2138 | 62.86 | 1244.82 | 14.55 | 264.72 | 22.59 | 374.41 | 1883.95 | 100/100 |
| **64** | 61.33 | 1388 | 1.68 | 38 | 1.1 | 25 | 35.88 | 812 | 2263 | 63.21 | 914.88 | 20.31 | 225.14 | 16.48 | 208.19 | 1348.21 | 100/100 |
| **65P** | 83.29 | 1730 | 0.19 | 4 | 0.29 | 6 | 16.23 | 337 | 2077 | 92.32 | 1334.67 | 3.41 | 37.79 | 4.27 | 43.89 | 1416.35 | 100/100 |
| **66** | 27.78 | 474 | 11.66 | 199 | 12.25 | 209 | 48.3 | 824 | 1706 | 61.94 | 143.51 | 20.49 | 64 | 17.57 | 51.97 | 259.48 | 100/100 |
| **67** | 28.04 | 496 | 12.04 | 213 | 11.93 | 211 | 47.99 | 849 | 1769 | 58.04 | 352.61 | 18.95 | 117.61 | 23.01 | 134.46 | 604.68 | 100/100 |
| **68** | 22.62 | 307 | 19.68 | 267 | 18.2 | 247 | 39.5 | 536 | 1357 | 29.75 | 129.02 | 41.42 | 167.65 | 28.84 | 115.8 | 412.47 | 60.5/66 |
| **69** | 10.42 | 115 | 13.32 | 147 | 26.09 | 288 | 50.18 | 554 | 1104 | 22.53 | 51.82 | 23.56 | 46.64 | 52.92 | 80.89 | 179.35 | 72.8/72 |
| **70** | 15.08 | 68 | 9.09 | 41 | 9.76 | 44 | 66.08 | 298 | 451 | 39.81 | 45.36 | 26.78 | 30.49 | 30.41 | 39.7 | 115.55 | 99.9/100 |
| **71** | 49.38 | 79 | 8.13 | 13 | 5 | 8 | 37.5 | 60 | 160 | 46.63 | 24.41 | 25.26 | 25.47 | 26.11 | 11.04 | 60.92 | 99.9/76 |
| **72** | 0 | 0 | 27.27 | 3 | 18.18 | 2 | 54.55 | 6 | 11 | 27.08 | 0.48 | 21.26 | 0.53 | 31.66 | 0.48 | 1.49 | 69.1/36 |
| **73** | 24.68 | 609 | 14.59 | 360 | 18.4 | 454 | 42.34 | 1045 | 2468 | 33.3 | 455.28 | 31.18 | 405.91 | 34.52 | 432.8 | 1293.99 | 100/100 |
| **74K** | 65.89 | 1219 | 4.11 | 76 | 4.92 | 91 | 25.08 | 464 | 1850 | 55.17 | 1418.75 | 21.57 | 542.85 | 23.26 | 582.56 | 2544.16 | 100/100 |
| **75** | 93.48 | 1519 | 1.29 | 21 | 1.11 | 18 | 4.12 | 67 | 1625 | 97.05 | 5954 | 1.57 | 77.16 | 1.38 | 72.79 | 6103.95 | 100/100 |
| **76*** | 49.78 | 1111 | 3.54 | 79 | 4.44 | 99 | 42.25 | 943 | 2232 | 53.45 | 829.73 | 22.42 | 329.9 | 24.12 | 346.2 | 1505.83 | 100/100 |
| **77**** | 45.29 | 77 | 10 | 17 | 5.29 | 9 | 39.41 | 67 | 170 | 53.88 | 77.6 | 22.06 | 32.33 | 23.06 | 30.33 | 140.26 | 100/100 |
| **78** | 41.44 | 46 | 6.31 | 7 | 7.21 | 8 | 45.05 | 50 | 111 | 46.15 | 62.92 | 26.86 | 36.97 | 26.99 | 35.81 | 135.7 | 100/100 |
| **79** | 30 | 261 | 7.24 | 63 | 12.64 | 110 | 50.11 | 436 | 870 | 39.49 | 690.88 | 30.02 | 510.42 | 30.49 | 513.47 | 1714.77 | 100/100 |
| **80** | 18.62 | 213 | 20.1 | 230 | 16.96 | 194 | 44.32 | 507 | 1144 | 35.56 | 403.29 | 34.21 | 384.59 | 30.23 | 338.2 | 1126.08 | 100/100 |
| **81** | 60.45 | 405 | 3.43 | 23 | 3.13 | 21 | 32.99 | 221 | 670 | 69.94 | 394.51 | 15.15 | 87.39 | 14.91 | 87.72 | 569.62 | 100/100 |
| **82** | 29.42 | 188 | 10.33 | 66 | 27.7 | 177 | 32.55 | 208 | 639 | 39.95 | 187.66 | 22.95 | 102.36 | 36.1 | 155.5 | 445.52 | 100/95 |
| **83** | 64.5 | 674 | 8.8 | 92 | 8.71 | 91 | 17.99 | 188 | 1045 | 55.05 | 675.37 | 23.63 | 276.34 | 21.33 | 264.77 | 1216.48 | 100/100 |
| **84** | 90.07 | 689 | 3.4 | 26 | 2.48 | 19 | 4.05 | 31 | 765 | 94.77 | 1690.34 | 2.92 | 52.45 | 2.31 | 41.06 | 1783.85 | 100/100 |
| **85** | 27.91 | 96 | 19.19 | 66 | 14.53 | 50 | 38.37 | 132 | 344 | 43.08 | 150.57 | 33.56 | 116.66 | 23.36 | 79.84 | 347.07 | 99.8/93 |
| **86** | 45.28 | 72 | 10.06 | 16 | 10.69 | 17 | 33.96 | 54 | 159 | 62.57 | 134.38 | 22.18 | 47.28 | 15.24 | 32.71 | 214.37 | 100/100 |
| **87** | 52.11 | 420 | 4.09 | 33 | 5.33 | 43 | 38.46 | 310 | 806 | 60.73 | 537.48 | 19.57 | 176.2 | 19.7 | 175.72 | 889.4 | 100/100 |
| **88** | 35.05 | 211 | 19.1 | 115 | 20.1 | 121 | 25.75 | 155 | 602 | 32.96 | 399.42 | 30.67 | 375.65 | 36.37 | 422.32 | 1197.39 | 100/100 |
| **89** | 65.6 | 328 | 6.4 | 32 | 9.2 | 46 | 18.8 | 94 | 500 | 50.98 | 651.46 | 24.23 | 302.81 | 24.79 | 311.04 | 1265.31 | 100/100 |
| **90** | 63.43 | 137 | 7.41 | 16 | 17.59 | 38 | 11.57 | 25 | 216 | 47.43 | 301.43 | 22.26 | 142.96 | 30.3 | 189.86 | 634.25 | 100/100 |
| **91** | 89.59 | 198 | 5.88 | 13 | 0.9 | 2 | 3.62 | 8 | 221 | 93.67 | 1015.96 | 3.64 | 38.89 | 2.69 | 30.24 | 1085.09 | 100/100 |
| **92** | 64.37 | 952 | 8.45 | 125 | 5.61 | 83 | 21.57 | 319 | 1479 | 52.98 | 670.62 | 26.98 | 329.59 | 20.04 | 244.25 | 1244.46 | 100/100 |
| **93** | 73.25 | 638 | 8.73 | 76 | 8.04 | 70 | 9.99 | 87 | 871 | 69.43 | 992.62 | 15.44 | 219.32 | 15.13 | 213.95 | 1425.89 | 100/100 |
| **94** | 90.05 | 516 | 2.62 | 15 | 2.27 | 13 | 5.06 | 29 | 573 | 97.45 | 1122.64 | 1.78 | 19.05 | 0.77 | 7.88 | 1149.57 | 100/100 |
| **95Z** | 63.41 | 792 | 7.53 | 94 | 9.37 | 117 | 19.7 | 246 | 1249 | 56.31 | 899.2 | 20.12 | 311.28 | 23.57 | 355.88 | 1566.36 | 100/100 |
| **96** | 44.83 | 507 | 5.13 | 58 | 3.98 | 45 | 46.07 | 521 | 1131 | 64.44 | 328.21 | 19.8 | 91.29 | 15.76 | 74.85 | 494.35 | 100/100 |
| **97** | 57.83 | 569 | 7.72 | 76 | 7.32 | 72 | 27.13 | 267 | 984 | 78.84 | 395.33 | 10.56 | 47.44 | 10.6 | 43.15 | 485.92 | 100/100 |

*Node 76 is the common ancestor of species with courtship signaling.
**Node 77 is the common ancestor of species with bioluminescence.
P Node 65 is the common ancestor of *Photeros*
K Node 74 is the common ancestor of *Kornickeria*
M Node 63 is the common ancestor of *Maristella*
Z Node 95 is the common ancestor of Z-group

### II. Bayesian relaxed molecular clock analyses of divergence times

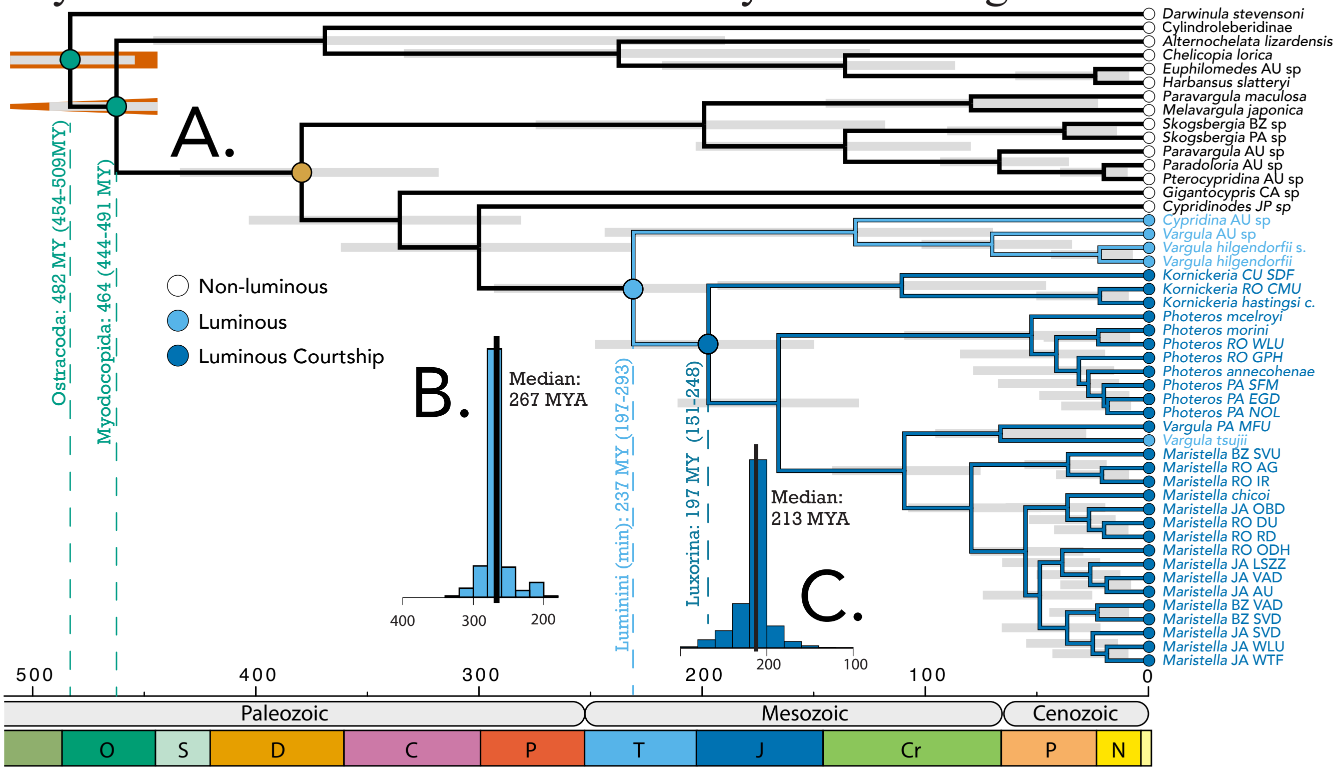


**Figure S5**. Relaxed molecular clock analysis. Figure 2 copied for more convenient comparison with Fig. S9. **A.** We used a Bayesian relaxed molecular clock approach, implemented in MrBayes (Ronquist and Huelsenbeck 2003) with 15 orthologs selected using sorta-date (Smith, Brown, and Walker 2018). We used two fossil constraints: Myodocopida, with offset exponential prior with a minimum of 448.8 MY and maximum of 509 MY and Ostracoda with a uniform prior between 443.8 and 509 MY (Oakley et al. 2013; Wolfe et al. 2016)  We used the Independent Gamma Rate (IGR) model in MrBayes, (Lepage et al. 2007; Chi Zhang 2016) with a prior of exp(10). We assumed a lognormal distribution with a mean of 0.001 and standard deviation of 0.0007 for the rate prior. We assumed fixed-rate amino acid models with aamodelpr=mixed. We ran 500K steps of Markov Chain Monte Carlo (MCMC) in two chains, using a starting phylogeny based on parsimony. **B.** Results of time-tree stochastic character mapping (ttscm) estimates origin of bioluminescence to be 267 MYA because that is the median age of character state transitions to bioluminescence **C.** Similarly, ttscm estimates the origin of luminous courtship to be 213 MYA.

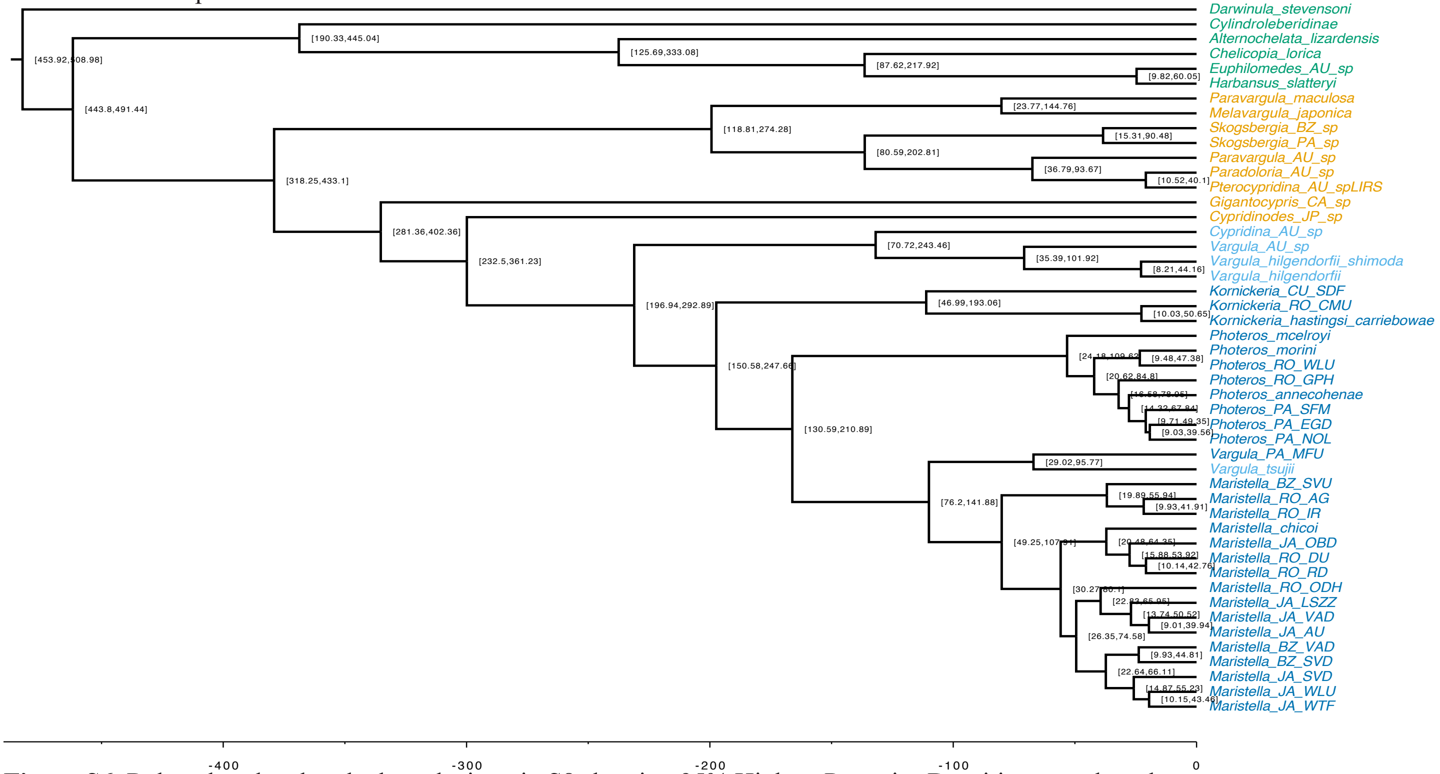


**Figure S6**. Relaxed molecular clock analysis as in S8 showing 95% Highest Posterior Densities at each node.

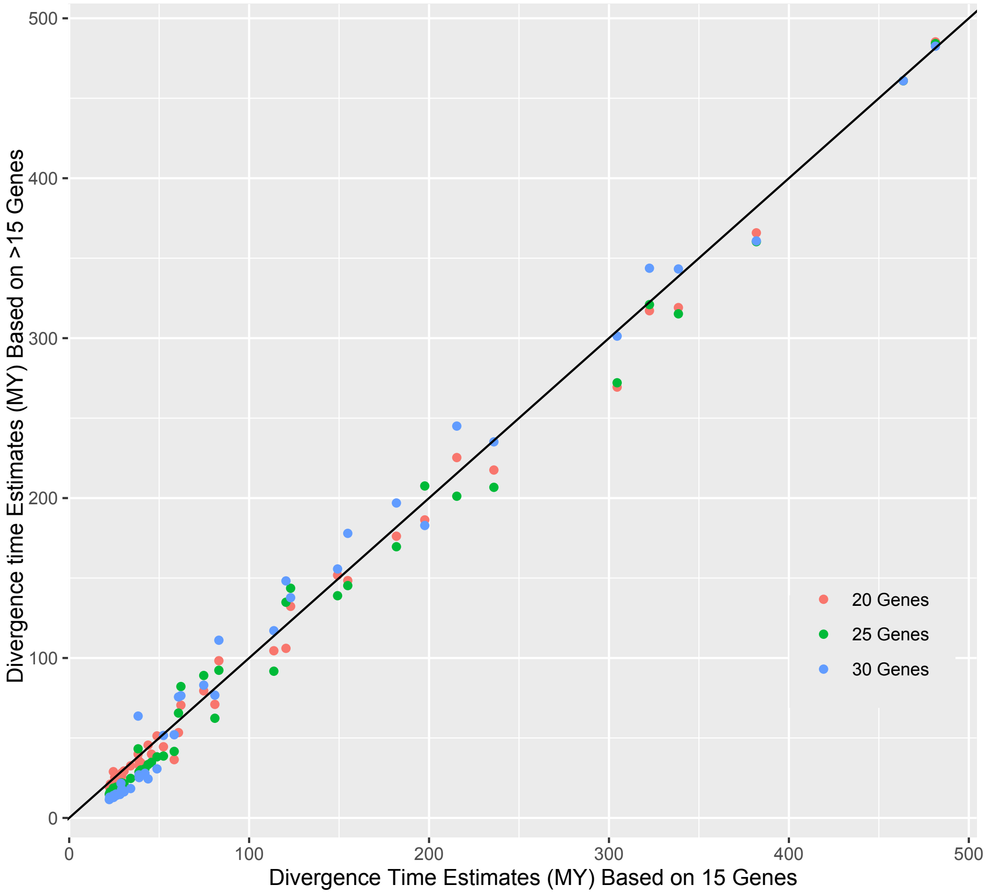


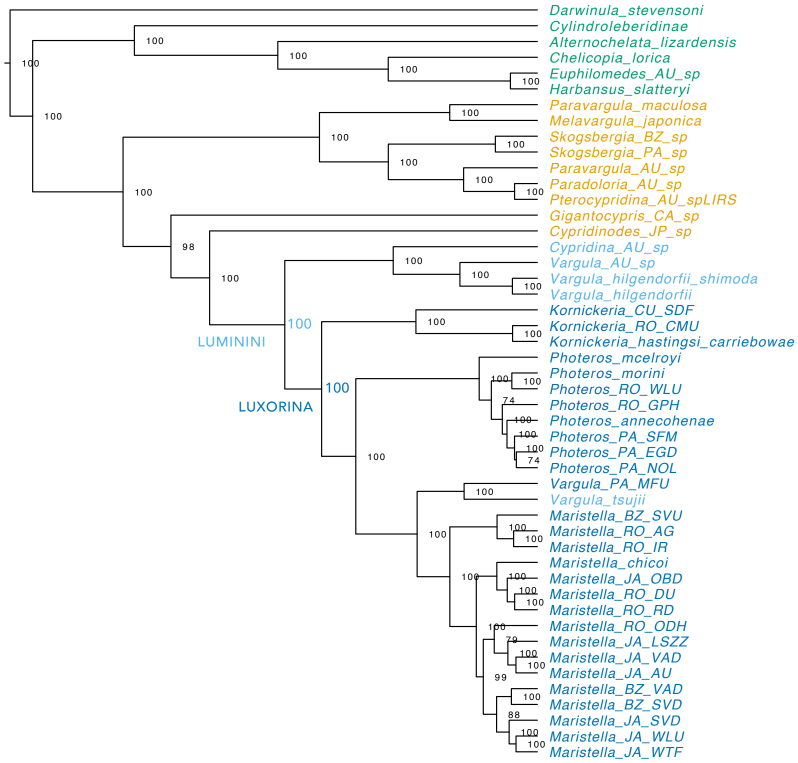


**Figure S7**. Sensitivity analysis of divergence time estimates using subsets of genes with Sortadate. We compared results from 15 (x-axis), 20, 25 and 30 (y-axis) genes, keeping all other parameters constant.

**Figure S8**. Posterior probability values of nodal support from Bayesian analysis using 15 genes selected with sortadate.

**Table S5**. Accession, sample, and collection data for mitochondrial data.

| Species (Fig S11) | Fig. 3 | Locale | 12S Accession | 16S Accession | CO1 Accession | Source | Synonymous sample names | Location | Collector | Year |
|---|---|---|---|---|---|---|---|---|---|---|
| Cylindroleberidinae | (not shown) | PP | N/A | DN177767_c165_g1_i1 | N/A | Schwentner et al (2018) | | | | |
| *Cypridinodes_sp* | Cypsp | PP | N/A | AY624732.1 | N/A | | | | | |
| *Gigantocypris_CA_sp* | Gigsp | CA | OL828721 | OL828505 | N/A | Torres and Gonzalez (2007) | Gigantocypris_sp | Santa Barbara, CA | S. Haddock | unknown |
| *Gigantocypris_CA_sp* | Gigsp | PP | N/A | N/A | KP745264.1 | Nigro et al (2016) | Gigantocypris_ muelleri | | | |
| *Paradoloria_pellucida* | Ppel | PP | N/A | AY624734.1 | N/A | Wakayama and Abe (2006) | | | | |
| *Paravargula_maculosa* | Pmac | AU | DN5047_c0_g1_i1 | N/A | DN6496_c0_g1_i1 | Transcriptome (Table S1) | BothPara_w | | | |
| *Heterodesmus_adamsii* | Hada | PP | N/A | MH784581.1 | MH815006.1 | Pham et al (2020) | | | | |
| *Skogsbergia_BZ_sp* | Skobz | BZ | DN16047_c0_g2 | N/A | N/A | Transcriptome (Table S1) | GlovSkog | | | |
| *Paradoloria_AU_sp* | Prvsp | AU | DN39598_c1_g2_i1 | N/A | DN32407_c0_g1_i1 | Transcriptome (Table S1) | FatWhiteEye | | | |
| *Pterocypridina_AU_ spLIRS* | Ptesp | AU | DN45931_c1_g6_i7 | N/A | N/A | Transcriptome (Table S1) | WhiteVanessa | | | |
| *Skogsbergia_lerneri* | Sler | BA | OL828722 | OL828506 | OL810011 | Torres and Gonzalez (2007) | Lerneri2017B | Bimini Id., Bahamas | E. Torres, V. Gonzalez | unknown |
| *Skogsbergia_crenulata* | Scre | VI | N/A | OL828507 | OL810012 | Torres and Gonzalez (2007) | crenulataUSVI01 | St. Thomas, US Virgin Islands | E.Torres | unknown |
| *Paradoloria_sp* | Parsp | AU | OL828723 | OL828508 | OL810013 | Torres and Gonzalez (2007) | Paradoloria_sp. VG | Jervis Bay, NSW, Australia | A. Parker | unknown |
| *Melavargula_sp* | Melsp | | N/A | OL828509 | N/A | Torres and Gonzalez (2007) | | Okinawa, Japan | M. Grygier | unknown |
| *Melavargula_japonica* | Mjap | JP | N/A | AY624733.1 | N/A | Transcriptome (Table S1) | | | | |
| *Vargula_norvegica* | Vnor | AT | OL828724 | OL828510 | N/A | Torres and Gonzalez (2007) | Vnorvegica | deep sea Atlantic sampling | A. Heger | unknown |
| *Vargula_karamu* | Vkar | AU | OL828725 | OL828511 | OL810014 | Torres and Gonzalez (2007) | Vargula_karamu = VarAP8 | Botany Bay, NSW, Australia | A. Parker | unknown |
| *Sheina_orri* | Sorr | AU | N/A | OL828512 | N/A | Torres and Gonzalez (2007) | Sheina_sp | Lizard Id., Australia | A. Parker | unknown |
| *Jimmorinia* | Jim | VI | OL828726 | OL828513 | N/A | Torres and Gonzalez (2007) | Jimmorinia | St. Thomas, US Virgin Islands | E. Torres | unknown |
| *Vargula_tubulata* | Vtub | AU | N/A | OL828514 | N/A | Torres and Gonzalez (2007) | Vtubulata | Jervis Bay, NSW, Australia | A. Parker | unknown |
| *Vargula_hilgendorfii* | Vhil | JP | NC_005306.1 | NC_005306.1 | NC_005306.1 | Ogoh and Ohmiya (2004) mt genome | Vargula hilgendorfii | | | |
| *Vargula_hilgendorfii_ shimoda* | Vhils | JP | N/A | N/A | DN8252_c1_g4_i1 | Transcriptome (Table S1) | | | | |
| *Cypridina_dentata* | Cden | PP | NC_042792.1 | NC_042792.1 | NC_042792.1 | Wang et al (2019) mt genome | | | | |
| *Cypridina_AU_spVG* | Cyp-spVG | AU | OL828727 | OL828515 | OL810015 | Torres and Gonzalez (2007) | Cypridina_sp = Cyp2A | Palm Beach, NSW, Australia | A. Parker | unknown |

Table S5 (continued). Sample and collection data for mitochondrial data

| Species (Fig S11) | Fig. 3 | Locale | 12S Accession | 16S Accession | CO1 Accession | Source | Synonymous sample names | Location | Collector | Year |
|---|---|---|---|---|---|---|---|---|---|---|
| *Cypridina_noctiluca* | Cnoc | JP | N/A | AY624731.1 | N/A | Wakayama and Abe (2006) | | | | |
| *Kornickeria_CU_SDF* | SDF | CU | DN58992_c0_g2_i1 | N/A | DN71800_c56_g6_i2 | Transcriptome (Table S1) | | | | |
| *Kornickeria_FL_GU* | GU | FL | OL828728 | OL828516 | OL810016 | Torres and Gonzalez (2007) | KornGU = Korn-GUB | Looe Key, Florida | E. Torres and J. Morin | 1992 |
| *Kornickeria_marleyi* | Kmar | JA | OL828729 | OL828517 | OL810017 | Torres and Gonzalez (2007) | Kornmarleyi = Mar30B | Discovery Bay, Jamaica | P. Edmunds | unknown |
| *Kornickeria_hastingsi* | Khas | BZ | OL828730 | OL828518 | OL810018 | Torres and Gonzalez (2007) | KhastingsiBZ0604D = BZ0706.04C | Carrie Bow Caye, Belize | E. Torres and A. Cohen | unknown |
| *Kornickeria_RO_CMU* | CMU | RO | N/A | DN224514_c3_g2_i2 | DN224514_c3_g2_i2 | Transcriptome (Table S1) | | | | |
| *Photeros_morini* | Pmor | BZ | DN109008_c0_g1_i2 | N/A | N/A | Torres and Gonzalez (2007) | PVFF = BZ1024A | Carrie Bow Caye, Belize | E. Torres and A. Cohen | unknown |
| *Photeros_graminicola* | Pgra | PA | OL828731 | OL828519 | OL810019 | Torres and Gonzalez (2007) | Pgraminicola = GRAMB | San Blas Islands, Panama | Jim Morin | 1993 |
| *Photeros_FL_WLOU* | WLOU | FL | OL828732 | OL828520 | N/A | Torres and Gonzalez (2007) | PWLOU = WlousA | Looe Key, Florida | E. Torres and J. Morin | 1992 |
| *Photeros_FL_QF* | QF | FL | OL828733 | OL828521 | OL810020 | Torres and Gonzalez (2007) | PQF = QFA | Looe Key, Florida | E. Torres and J. Morin | 1992 |
| *Photeros_shulmanae* | Pshu | PA | N/A | OL828522 | N/A | Torres and Gonzalez (2007) | Pshulmanae = Pshul | San Blas Islands, Panama | Jim Morin | 1993 |
| *Photeros_annecohenae* | Pann | BZ | DN86176_c0_g3_i1 | N/A | N/A | Hensley et al 2020 Transcriptome | Gruber_Pannecohenae | | | |
| *Photeros_PA_SFM* | SFM | PA | DN21_c0_g2_i2 | DN21_c0_g2_i2 | DN21_c0_g2_i2 | Transcriptome (Table S1) | | | | |
| *Photeros_PA_EGD* | EGD | PA | DN488_c0_g2_i2 | N/A | DN488_c0_g2_i2 | Transcriptome (Table S1) | | | | |
| *Photeros_PA_NOL* | NOL | PA | DN760_c0_g1_i10 | DN760_c0_g1_i10 | DN760_c0_g1_i10 | Transcriptome (Table S1) | | | | |
| *Vargula_tsujii* | Vtsu | CA | NC_039175.1 | NC_039175.1 | NC_039175.1 | Goodheart et al (2020) mt genome | | | | |
| *Vargula_PA_MFU* | MFU | PA | DN9711_c0_g1_i1 | N/A | N/A | Transcriptome (Table S1) | | | | |
| *Vargula_mizonomma* | Vmiz | PA | OL828734 | OL828523 | OL810021 | Torres and Gonzalez (2007) | ZGRPmizonomma = VMizoCD | San Blas Islands, Panama | Jim Morin | 1993 |
| *Vargula_kuna* | Vkun | PA | OL828735 | OL828524 | N/A | Torres and Gonzalez (2007) | kuna1B = Kuna934A | San Blas Islands, Panama | Jim Morin | 1993 |
| *Maristella_chicoi* | Mchi | BZ | N/A | N/A | N/A | Transcriptome (Table S1) | | | | |
| *Maristella_JA_LSZZ* | LSZZ | JA | DN182361_c2_g1_i5 | N/A | N/A | Transcriptome (Table S1) | | | | |
| *Maristella_JA_AU* | AU | AU | DN57492_c0_g2_i1 | N/A | DN56818_c0_g1_i1 | Transcriptome (Table S1) | | | | |
| *Maristella_JA_VAD* | JVAD | JA | N/A | N/A | DN94762_c1_g2_i2 | Transcriptome (Table S1) | | | | |
| *Maristella_BZ_VAD* | BVAD | BZ | DN209300_c0_g3_i2 | DN209300_c0_g3_i2 | N/A | Transcriptome (Table S1) | | | | |

Table S5 (continued). Sample and collection data for mitochondrial data

| Species (Fig S11) | Fig. 3 | Locale | 12S Accession | 16S Accession | CO1 Accession | Source | Synonymous sample names | Location | Collector | Year |
|---|---|---|---|---|---|---|---|---|---|---|
| *Maristella_BZ_SVD* | BSVD | BZ | N/A | N/A | DN25384_c0_g1_i2 | Transcriptome (Table S1) | BZZZDB | South Water Caye, Belize | G. Gerrish | unknown |
| *Maristella_JA_WLU* | JWLU | JA | DN60033_c11_g1_i1 | N/A | DN60033_c11_g1_i1 | Transcriptome (Table S1) | | | | |
| *Maristella_JA_OBD* | JOBD | JA | DN43094_c0_g1_i1 | DN43094_c0_g1_i1 | N/A | Transcriptome (Table S1) | | | | |
| *Maristella_RO_RD* | RD | RO | DN62759_c0_g4_i1 | N/A | DN58584_c0_g1_i1 | Transcriptome (Table S1) | | | | |
| *Maristella_JA_SVD* | JSVD | JA | N/A | N/A | DN43495_c0_g2_i1 | Transcriptome (Table S1) | | | | |
| *Maristella_RO_ODH* | ODH | RO | DN54797_c3_g3_i3 | N/A | DN13681_c0_g1_i1 | Transcriptome (Table S1) | | | | |
| *Vargula_FL_CGrp* | CGrp | FL | OL828736 | N/A | OL810022 | Torres and Gonzalez (2007) | CGRPFlorida = HC81B | Looe Key, Florida | E. Torres and J. Morin | unknown |
| *Vargula_contragula* | Vcon | PA | N/A | OL828525 | OL810023 | Torres and Gonzalez (2007) | CGRPcontragula = Contragula | San Blas Islands, Panama | Jim Morin | 1993 |
| *Maristella_micamacula* | Mmic | PA | OL828737 | OL828526 | OL810024 | Torres and Gonzalez (2007) | UGRP_micamacula = VMicaC | San Blas Islands, Panama | Jim Morin | 1993 |
| *Maristella_scintilla* | Msci | PA | OL828738 | OL828527 | OL810025 | Torres and Gonzalez (2007) | RGRP_scintilla = VSCINB | San Blas Islands, Panama | Jim Morin | 1993 |
| *Maristella_BZ_SVU* | BSVU | BZ | DN29269_c4_g2_i3 | N/A | N/A | Transcriptome (Table S1) | RGRPMWU_BZ0313 = BZ0313A | Carrie Bow Caye, Belize | E. Torres and A. Cohen | unknown |
| *Maristella_noropsela* | Mnor | BZ | OL828739 | OL828528 | OL810026 | Torres and Gonzalez (2007) | RGRPnoropsela = VNorop17 | San Blas Islands, Panama | Jim Morin | 1993 |
| *Maristella_RO_AG* | AG | RO | DN382053_c0_g2_i1 | DN382053_c0_g2_i1 | DN382053_c0_g2_i1 | Transcriptome (Table S1) | | | | |
| *Maristella_RO_IR* | IR | RO | N/A | N/A | DN36910_c5_g1_i1 | Transcriptome (Table S1) | | | | |

### III. Expanded taxon sampling with mitochondrial genes

**Figure S11**. Copy of Figure 3. Expanded taxon sampling by adding individual mitochondrial genes, mainly from Torres and Gonzalez (2007) to the transcriptome data in Figure 1. We inferred this species phylogeny using the multi-species-coalescent, employed with ASTRAL-PRO (Chao Zhang et al. 2020), treating multiple mitochondrial genes as a single locus. Taxa colored pink are those that contain only mitochondrial data. Locality abbreviations are as in Fig. 1 and also include additional locations of VI=US Virgin Islands and FL=Florida USA. For *Jimmorinia* (*), we combined data from transcriptomes from a Jamaican species (cf *J. gunnari*) with mitochondrial DNA from a specimen (*Jimmorinia* sp USVI) unidentified to species from VI. We included outgroups in the analysis, but they are not illustrated here.

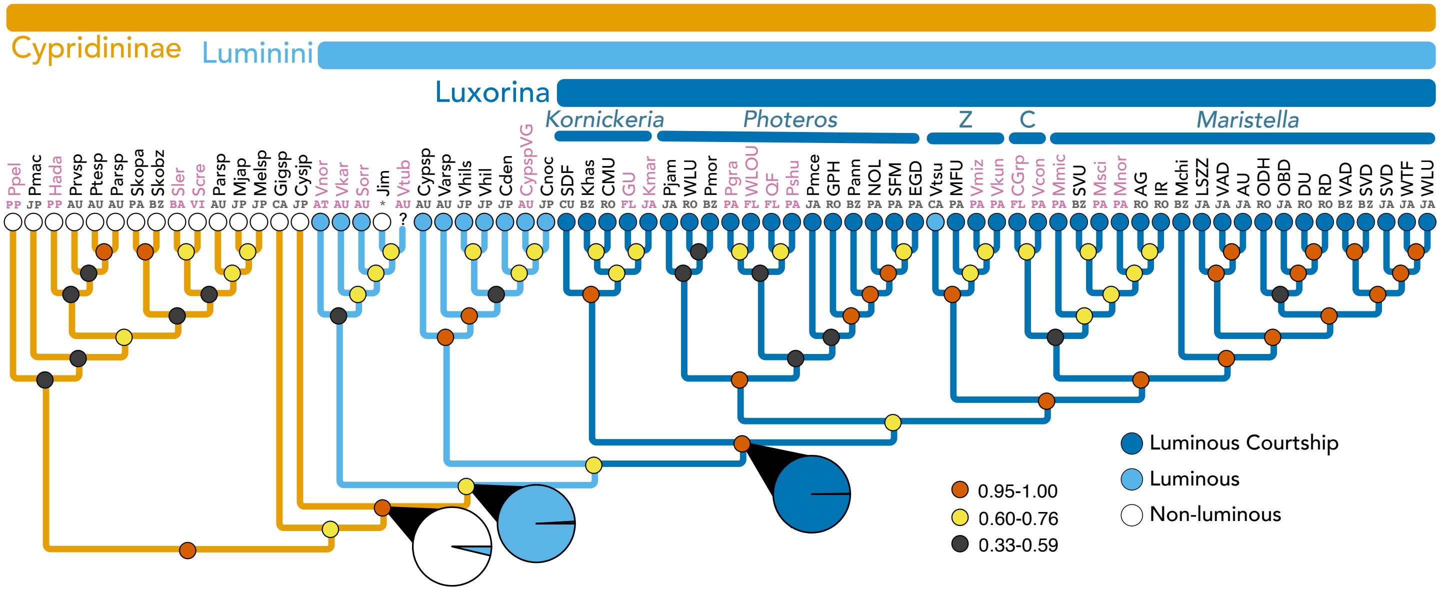


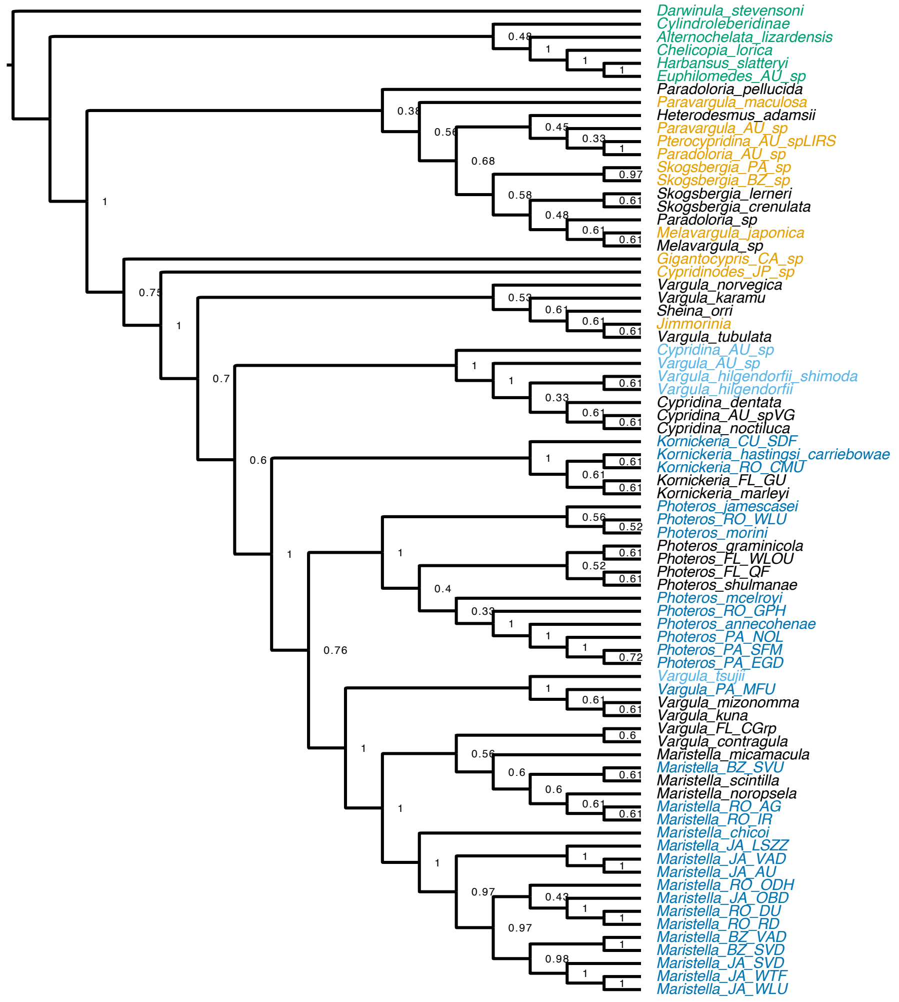


**Figure S9**. Multi-species coalescent species tree (equivalent to Figs 3, S11) combining transcriptome data with previously published mitochondrial data. This figure has full species names rather than abbreviations. Names colored black have only mitochondrial data. Numbers at nodes are posterior probability values. See Figs 3 and S9 for detailed methods.

**Figure S10**. Maximum likelihood phylogeny based on mitochondrial data alone, including 12S, 16S, and co1. Taxa in colors use mitochondrial data from transcriptomes. We found in each transcriptome, the longest contigs with high similarity to published mitochondrial genomes of cypridinid ostracods. We then annotated the contigs using MITOS and concatenated 12S, 16S, and co1 data with that of Torres and Gonzalez (2007) and GenBank data (see Table S5). We assumed best-fit models in IQ-TREE and searched for the ML tree. Numbers at nodes are ultrafast bootstrap and alrt support based on 1000 replicates each.

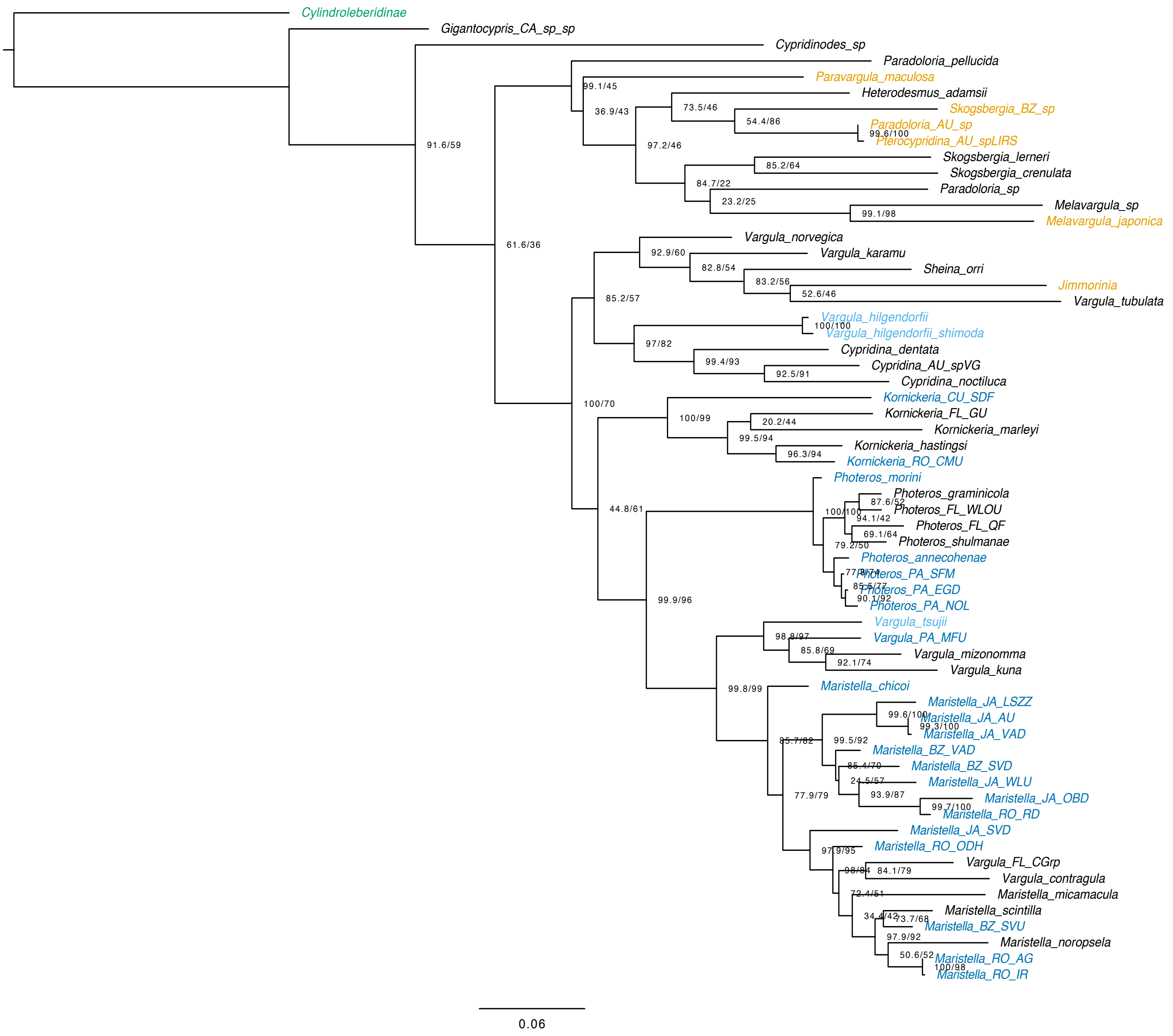

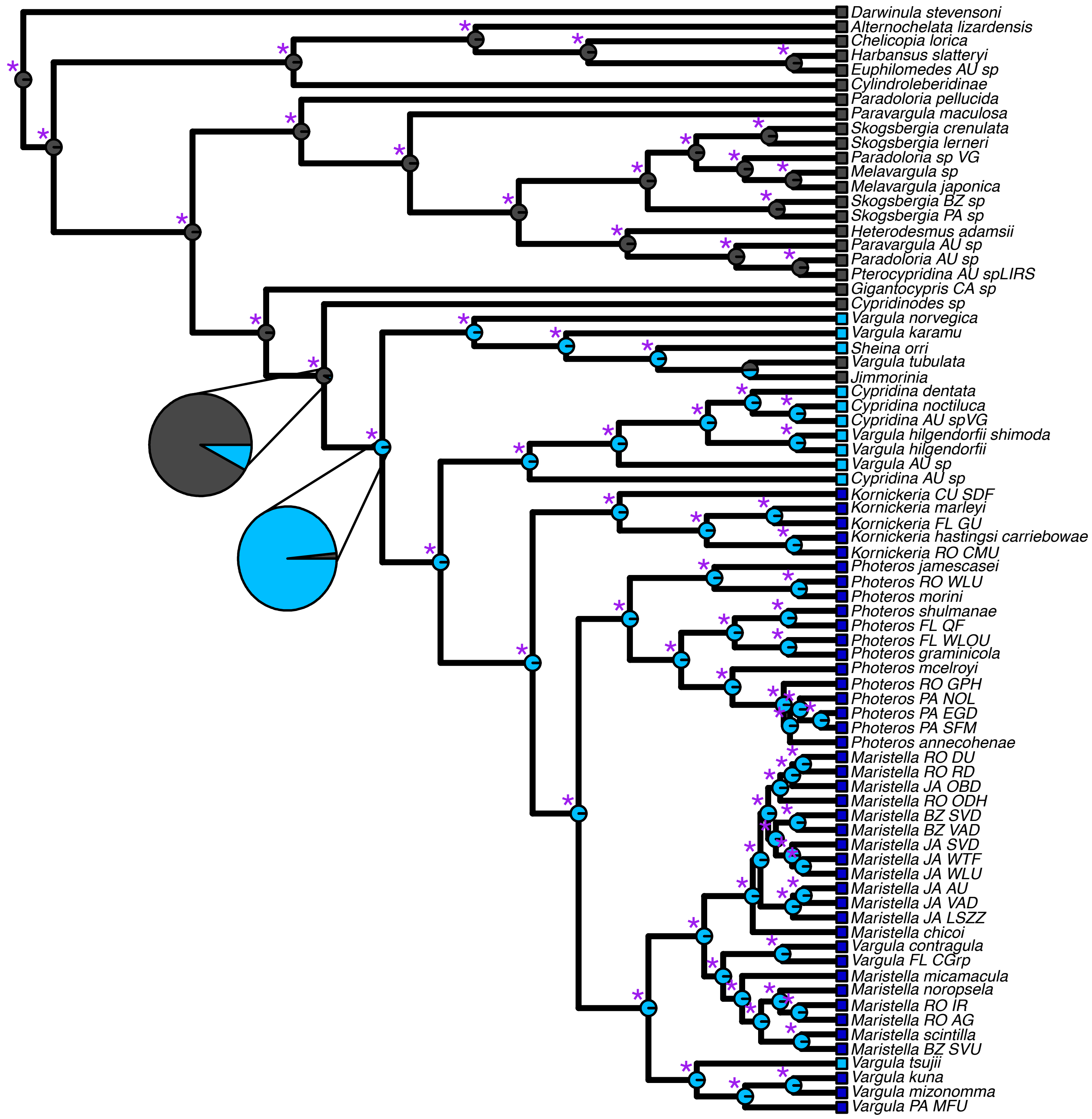


**Figure S11**. Ancestral state reconstruction supports a single origin of bioluminescence in Cypridinidae. Ancestral state reconstruction analysis for the presence (blue) or absence (grey) of bioluminescence. Pie charts on the nodes are scaled marginal likelihoods calculated using the ace function in APE. The asterisks denote significant nodes as defined by proportional likelihood significance tests (Pagel 1999) with a likelihood difference of at least 2 or higher.

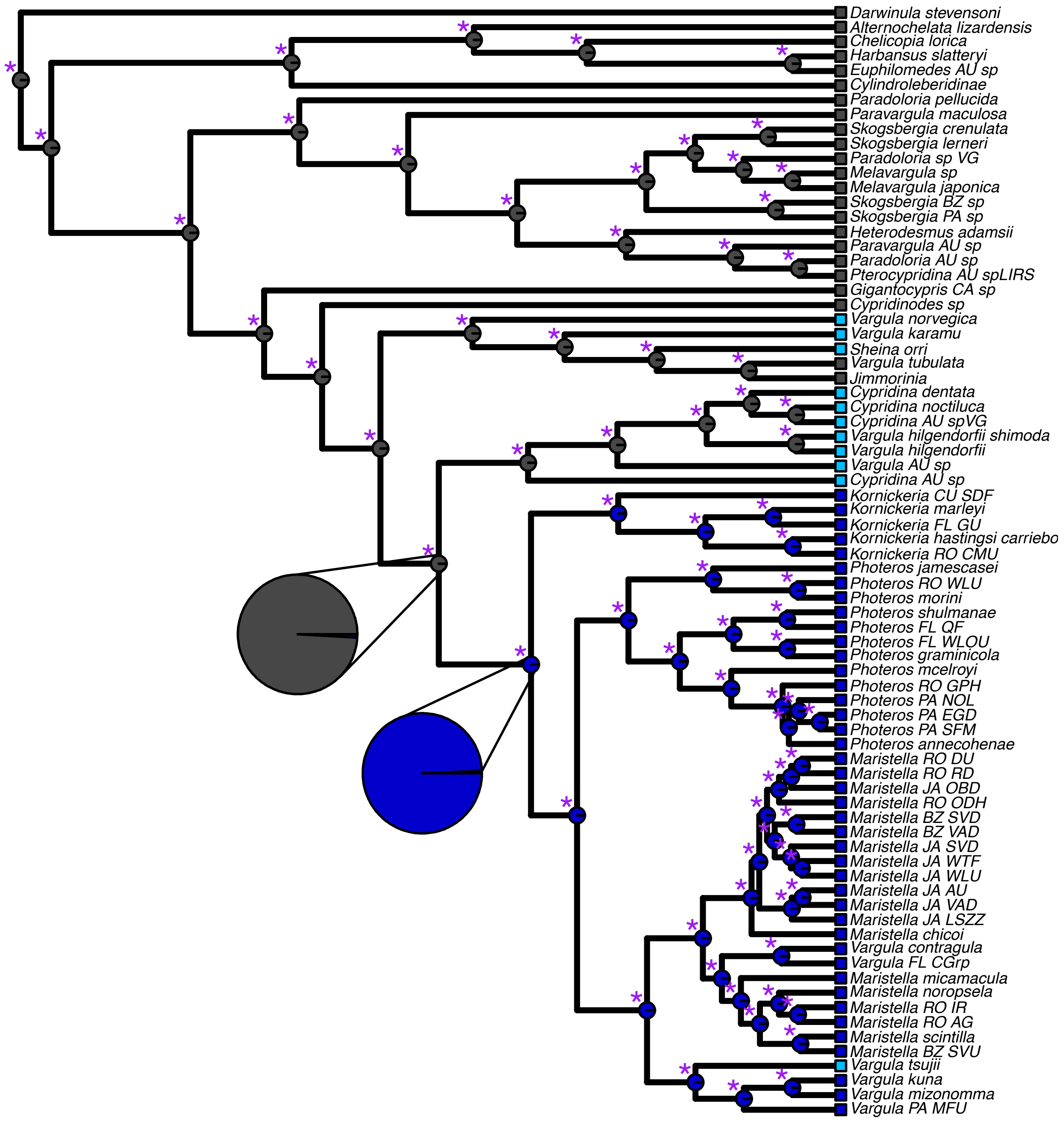


**Figure S12**. Ancestral state reconstruction supports a single origin of bioluminescent courtship in Cypridinidae. Ancestral state reconstruction analysis for the presence (blue) or absence (grey) of bioluminescent courtship displays. Pie charts on the nodes are scaled marginal likelihoods calculated using the ace function in APE. The asterisks denote significant nodes as defined by proportional likelihood significance tests (Pagel 1999) with a likelihood difference of at least 2 or higher.
